# CRISPR-Cas12a-assisted PCR tagging of mammalian genes

**DOI:** 10.1101/473876

**Authors:** Julia Fueller, Konrad Herbst, Matthias Meurer, Krisztina Gubicza, Bahtiyar Kurtulmus, Julia D. Knopf, Daniel Kirrmaier, Benjamin C. Buchmuller, Gislene Pereira, Marius K. Lemberg, Michael Knop

**Author notes:** These authors contributed equally to this work (Julia Fueller, Konrad Herbst, Matthias Meurer). These authors contributed equally to this work (Krisztina Gubicza, Bahtiyar Kurtulmus).

## Abstract

Here we describe a time-efficient strategy for endogenous C-terminal gene tagging in mammalian tissue culture cells. An online platform is used to design two long gene-specific oligonucleotides for PCR with generic template cassettes to create linear dsDNA donors, termed PCR cassettes. PCR cassettes encode the tag (e.g. GFP), a Cas12a CRISPR RNA for cleavage of the target locus and short homology arms for directed integration via homologous recombination. The integrated tag is coupled to a generic terminator shielding the tagged gene from the co-inserted auxiliary sequences. Co-transfection of PCR cassettes with a Cas12a-encoding plasmid leads to robust endogenous expression of tagged genes, with tagging efficiency of up to 20% without selection, and up to 60% when selection markers are used. We used target-enrichment sequencing to investigate all potential sources of artefacts. Our work outlines a quick strategy particularly suitable for exploratory studies using endogenous expression of fluorescent protein tagged genes

## Introduction

Targeted insertions of transgenes into genomes of mammalian cells (‘knock-ins’) for applications such as protein-tagging are critical genomic modifications for functional studies of genes within their endogenous context, thus reducing the likelihood of artifacts due to overexpression (Doyon et al., 2011). In mammalian cells such knock-ins are complicated by inefficient targeting and a high likelihood of off-target integrations. Knock-in efficiency in mammalian cells can be enhanced by inducing site-specific double strand breaks (DSBs) using programmable endonucleases such as zinc finger nucleases, transcription activator-like effector nucleases (TALENs) or clustered regularly interspaced short-palindromic repeat (CRISPR)-associated (Cas) endonucleases (Dambournet et al., 2014). These lesions can promote integration of the desired heterologous DNA sequences via DSB repair pathways such as homologous recombination (HR) or canonical non-homologous end joining (c-NHEJ) (Scully et al., 2019). While zinc finger nucleases and TALENs where initially shown to yield high on-target editing rates, the CRISPR-Cas endonucleases are nowadays preferred due to their simplistic usage and versatility (Zhang, 2019). Among different Cas-endonucleases, Cas9 has found its way into most genome engineering applications mainly because for historical reasons. The subsequently characterized Cas12a in comparison has the reported advantage of being more specific *in vivo* (Kleinstiver et al., 2016; Kim et al., 2017) and its CRISPR RNA (crRNA) structure is simpler (Zetsche et al., 2015). In addition, Cas9 induces DSBs close to the PAM site while Cas12a cuts further away from it, which might increase targeting efficiency as the target sequence is not as easily destroyed by indel formation and may be re-cleaved after repair (Zetsche et al., 2015; Moreno-Mateos et al., 2017).

A variety of methods have been developed to use Cas9/12a for knock-in applications (reviewed in (Yamamoto and Gerbi, 2018)). They can be classified by the DSB repair pathway they depend on. Methods that rely on c-NHEJ require a correctly positioned cut site for the endonuclease and alternative processing of the DNA ends can generate out of frame integrations. Methods relying on HR are more flexible in terms of target sites enabling highly precise genomic modifications. However, HR is only active in late S/G2 phase of the cell cycle (Moynahan and Jasin, 2010) decreasing the likelihood that this pathway is selected for the repair of a particular DSB. Irrespective of the method and targeted DNA repair pathway, suitable reagents are required to provide all the necessary components for integration such as recombinant proteins, RNAs, single-stranded DNA or the cloning of tailored and gene-specific plasmids (Yamamoto and Gerbi, 2018). In yeast genomic tagging has been simplified to a strategy based on PCR (Baudin et al., 1993; Wach et al., 1994), now commonly referred to as ‘PCR tagging’. It requires two gene-specific DNA oligonucleotides (oligos) for PCR and a generic ‘template plasmid’ that provides the tag and a selection marker to generate a ‘PCR cassette’. Upon transformation into cells the homologous sequences provided by the oligos target precise insertion of the PCR cassette into the genome by the efficient HR machinery in this species.

In mammalian cells long linear double stranded DNA donors containing short homology arms (50-100 bp) have been shown to suffice for efficient HR if a DSB is simultaneously induced at the modification site (Orlando et al., 2010; Zheng et al., 2014; Zhang et al., 2017). Hence, the use of PCR for the generation of repair templates for gene tagging in mammalian cells is in principle possible. However, it is complicated by the requirement to simultaneously introduce a crRNA for CRISPR-Cas-mediated cleavage at the modification site.

Here we develop ‘mammalian PCR tagging’. Similar to yeast PCR tagging, this method also depends on two gene-specific oligos and a single PCR. In contrast to the yeast method one of the oligos also contains the sequences encoding the crRNA and the PCR generates a fragment termed ‘PCR cassette’ that simultaneously contains a functional gene for the expression of the crRNA to direct the integration of the cassette into the genome. We optimized the design of the oligos and explored the effect of oligo protection by chemical modification, the use of selection markers and applications in different cell lines. Using targeted next generation sequencing we characterize tagging fidelity, off-target insertions and by-product formation such as repair template concatemerization. We facilitate adaptation of mammalian PCR tagging by introducing a toolbox comprising many possible PCR templates allowing genomic integration of various different tags. A web application allows the rapid design of the two oligos needed for mammalian PCR tagging of individual genes. Finally, we discuss applications of the method for basic research in cell biology and for screening purposes.

## Results

### Implementation of mammalian PCR tagging and method optimization

Mammalian PCR tagging requires two oligos for C-terminal tagging of proteins. The M1 tagging oligo provides homology to the 5’ region of the insertion site. The M2 tagging oligo provides homology to the 3’ region of the insertion site as well as the sequence of the crRNA for guiding the Cas12a endonuclease along with a (T)_6_ element that functions as a Pol III terminator (Arimbasseri et al., 2013). The PCR with the M1/M2 tagging oligos is performed with the template plasmid, which provides the desired tag (e.g. GFP) and a U6 polymerase III (Pol III) promoter for the crRNA. The template plasmid contains also a heterologous 3’-untranslated region (UTR) after the fluorescent protein reporter to properly terminate the gene fusion before the crRNA expression unit. PCR generates a ‘PCR cassette’ that contains locus-specific homology arms as well as a functional gene for the expression of a locus specific crRNA for Cas12a (Fig. 1a, S1). Based on our experience with similar PCR cassettes in yeast (Buchmuller et al., 2019) we predicted that upon transfection the crRNA will be expressed and will assemble with Cas12a, which is simultaneously expressed from a co-transfected plasmid, into a functional complex that cleaves the target gene (Fig. 1a,b).

**Figure 1.**
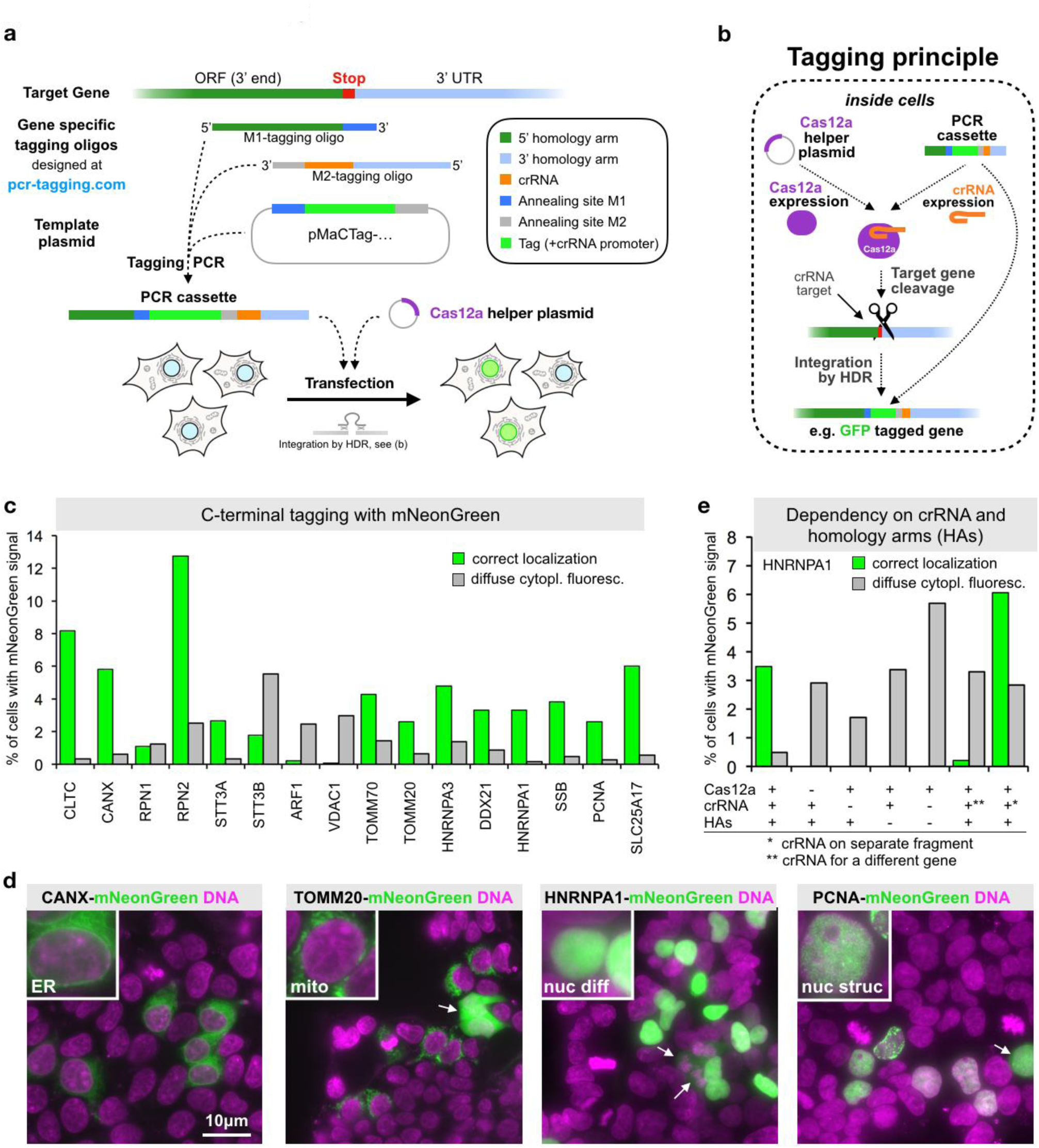
Endogenous C-terminal gene tagging in mammalian cells using PCR tagging. (**a**) For tag insertion before the STOP codon of an ORF, two ‘Gene specific tagging oligos’ (termed M1 and M2) are designed using an online tool (www.pcr-tagging.com). A ‘Tagging PCR’ with a generic ‘Template plasmid’ generates the gene-specific ‘PCR cassette’. The ‘Template plasmid’ provides the tag (e.g. a fluorescent protein), a possible selection marker and a Pol III promoter. For gene tagging the ‘PCR cassette’ is transfected into the target cell together with a helper plasmid containing a Cas12a endonuclease gene. This leads to insertion of the PCR cassette into the chromosome, which yields a fusion of the tag (e.g. GFP) with the target gene. (**b**) Tagging Principle: The PCR cassette contains a crRNA sequence that is expressed inside the cell via an U6 promoter (Pol III promoter). The crRNA directs Cas12a (which is expressed from the helper plasmid) to the target locus close to the insertion site. Stimulated by the DSB the linear PCR cassette is then inserted into the genome. The homology arm of the M1 tagging oligo thereby directs in frame fusion of the tag with the target ORF, leading to the expression of a tagged protein from the target locus. Integration leads to destruction of the crRNA target site, thus preventing re-cleavage of the modified locus. (**c**) Efficiency of C-terminal mNeonGreen-tagging for 16 organelle specific genes. For each gene, specific M1/M2 tagging oligos were used to amplify an mNeonGreen containing template plasmid. The resulting PCR cassettes were transfected in HEK293T cells. HOECHST staining of live cells and analysis by fluorescence microscopy was performed three days after transfection. Fractions of cells exhibiting the expected localization or diffuse cytoplasmic green fluorescence are shown. For information on selected genes, see Table S1. Data from one representative experiment is shown. (**d**) Representative images from HEK293T cells 3 days after transfection. mNeonGreen fluorescence and HOECHST staining (DNA) is shown. In addition to the expected localization, cells showing diffuse cytoplasmic fluorescence (arrows) are detected. (**e**) Tagging is specific for the crRNA and guided by the homology arms (HAs). Efficiency of control transfections (see Fig. S2 for representative examples). * in this transfection indicates that a matching combination of crRNA and HAs was used, but the crRNA was expressed from a different PCR fragment. ** indicates that in this case a PCR cassette was used where the crRNA (for CANX) lead to cleavage of a different gene than the one specified by the HAs (HNRNPA1). A small fraction of cells (<0.02%, corresponding to five cells in the entire well) exhibiting an ER localization pattern typically seen for CANX was observed, indicating cassette integration at the CANX locus, e.g. via c-NHEJ.

DSB repair can occur via different pathways. One option is that the DSB is repaired by HR using the transfected PCR cassette as template, as it contains homology arms that match the region adjacent to the cleaved site. This yields the desired integrands expressing the appropriately tagged proteins from the target locus. Other repair pathways like c-NHEJ are less well defined and likely do not produce a functionally tagged gene.

To test if this approach permits efficient gene tagging in mammalian cells we designed a template plasmid containing the bright green fluorescent protein mNeonGreen (Shaner et al., 2013). We designed 16 M1/M2 tagging oligo pairs for tagging of 16 different genes encoding proteins with a diverse range of cellular localizations (Table S1) and with high endogenous expression levels (Geiger et al., 2012; Schaab et al., 2012). This allows for easy detection of the corresponding mNeonGreen-tagged fusion proteins by fluorescence microscopy. We co-transfected the PCR cassettes together with a Cas12a encoding plasmid into HEK293T cells and quantified fluorescent cells three days later. For all genes, we observed between 0.2% and 13% of fluorescent cells with the expected protein-specific localization pattern (Fig. 1c), e.g. Endoplasmic Reticulum (ER) for CANX, mitochondrial staining for TOMM20, or a diffuse and a dotted nuclear staining for HNRNPA1 and PCNA, respectively (Fig. 1d). We validated that the formation of cells with correctly localized fluorescence signal depended on the presence of Cas12a and matching combinations of homology arms and crRNA, irrespective of whether they are on the same, or different PCR products (Fig. 1e). In the presence of a crRNA for a locus different from the one targeted by the homology arms, we found very rarely cells where the cassette became integrated into the foreign locus, indicating that in addition to HR also other integration pathways such as c-NHEJ are used (Fig. 1e and Fig. S2). Together, these results establish that the crRNA is transcribed from the transfected PCR cassette and that it directs Cas12a for cleavage of the target locus. Furthermore, we conclude that the Cas12a-mediated DSB is repaired frequently using HR and linear donor templates with short homology arms.

In addition to cells with the expected localization of the green fluorescence we observed in several transfections also cells with diffuse cytoplasmic fluorescence of variable brightness (Fig. 1c-d, see examples labeled with arrows in 1d). This fluorescence was independent on Cas12a or matching combinations of crRNA and homology arms (Fig. 1e). This indicates that the diffuse cytoplasmic signal resulted from the transfected PCR cassettes alone.

### Diffuse cytoplasmic fluorescence is caused by unstable extra-chromosomal DNA molecules

The nature of the diffuse cytoplasmic fluorescence observed in a fraction of the cells was unclear. We reasoned that the cytoplasmic fluorescence could originate from extra-chromosomal DNA molecules or fragments that have integrated at chromosomal off-target loci. To investigate the fate of the transfected fragments we specifically amplified from cells 3 days after transfection the junctions between PCR cassettes and their upstream flanking DNA sequences using Anchor-Seq (Meurer et al., 2018; Buchmuller et al., 2019) (Fig. 2a). We detected junctions indicative for PCR cassettes inserted into the correct chromosomal locus (Fig. 2b). However, the detection sensitivity of correctly inserted cassettes was limited because of a large number of reads that did not extend beyond the sequence of the M1 or M2 tagging oligos (Fig. 2b). This suggests that they result from transfected PCR cassettes that are still present in the cultured cells. In addition, we also observed a substantial fraction of reads that originate from ligated ends of transfected cassettes, consistent with the idea that the free ends of the transfected PCR cassettes were recognized and processed by c-NHEJ. Although different types of fusions were detected, the most dominating comprised a head-to-tail fusion of the PCR fragment (Fig. 2b).

**Figure 2.**
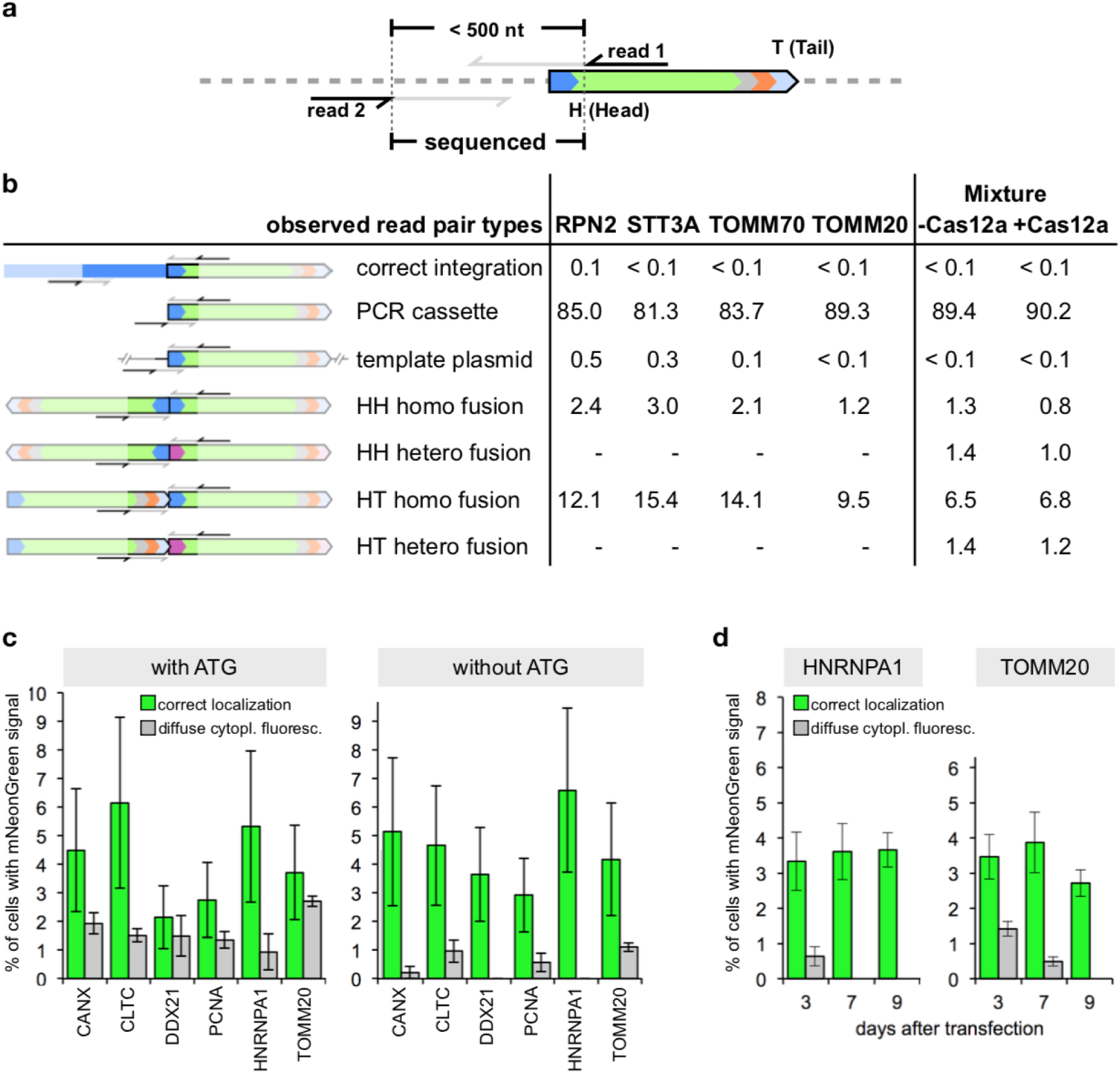
Analysis of the fate of the transfected PCR cassette using target enrichment sequencing. (**a**) Anchor-Seq (Meurer et al., 2018) is based on a target enrichment procedure that uses an oligo in the mNeonGreen gene to enrich adjacent sequences for analysis by next generation sequencing using a paired end sequencing protocol (reads 1 & 2). (**b**) Anchor-Seq analysis of adjacent sequences of the PCR cassette from HEK293T cells three days after transfection, for the four genes shown individually, and from cells transfected with a mixture of PCR cassettes for different genes (cassettes from the genes shown in Fig. 1c; labeled with ‘Mixture’). Fraction of reads (in %) observed for the different categories, where H and T stand for head and tail of the PCR cassette, respectively. Combinations of the letter denote the detected fusion, homo denotes fusion of two ends from a PCR cassette targeting the same gene, hetero from PCR cassettes targeting different genes. (**c**) HEK293T cells transfected with PCR cassettes as indicated using wild type mNeonGreen gene or lacking ATG translation initiation codons within the first 10 codons of the mNeonGreen ORF. Live cell fluorescence microscopy of HOECHST stained cells was used to determine the fraction of cells (in %) with correct localization and diffuse cytoplasmic fluorescence. Data from three replicates is shown. Error bars indicate SD. (**d**) HEK293T cells transfected with PCR cassettes for HNRNPA1 or TOMM20 were passaged for the indicated time periods. Analysis as in (c). Data from three replicates is shown. Error bars indicate SD.

To further explore the nature of the cassette fusions, we transfected a mixture of PCR cassettes used for tagging the genes shown in Fig. 1c. This detected hybrid-fusions between PCR cassettes targeting different genes (Fig. 2b), validating the idea that after transfection the cassettes are ligated together, e.g. via c-NHEJ mediated DNA damage repair. However, head-to-tail fusions among cassettes for the same gene remained the most abundant events also in the transfection of the mixture. This can be best explained by a preference for intramolecular ligation and subsequent concatemerization by HR, as reported in previous studies (Folger et al., 1982; 1985).

In head-to-tail fusions the crRNA gene is ligated to the 3’ end of the mNeonGreen sequence with the homology arms of the M1 and M2 tagging oligos in between. The used U6 Pol III promoter has previously been shown to also mediate Pol II driven expression (Rumi et al., 2006; Gao et al., 2018).

This could lead to the expression of mNeonGreen. To assess this, we next transfected a PCR cassette where the ATG codons at position 1 and 10 of the mNeonGreen open reading frame (ORF) have been substituted with a codon for valine, respectively. This largely, but not completely, suppressed the population of cells with diffuse cytoplasmic signal, while the fraction of cells with specific localization indicative for correct gene tagging was unchanged (Fig. 2c). This indicates that the necessary ATG is often provided by mNeonGreen itself. Additionally, the crRNA or homology sequences within the M1 or M2 tagging oligo may provide an ATG in frame with the mNeonGreen ORF.

If head-to-tail fused PCR cassettes are not or rarely incorporated into the genome they are unlikely to be stable. Consistently, we observed during subsequent growth of the cells a gradual loss of the fraction of cells with diffuse cytoplasmic fluorescence, while the fraction of cells with correctly localized fluorescence signal remained constant (Fig. 2d). This argues that head-to-tail fused fragments that are formed as byproducts, do not hamper a general applicability of mammalian PCR tagging for targeted ‘knock-in’ of PCR cassettes.

### Parameters influencing tagging efficiency

To explore PCR tagging further, we determined tagging efficiency as a function of various parameters.

#### DNA delivery

We first explored basic parameters such as DNA amount and transfection method. We found that equal amounts of Cas12a plasmid DNA and PCR cassette DNA are optimal (Fig. S3a), whereas the transfection method did not seem to influence the outcome (Fig. S3b). Furthermore, we noticed that PCR cassette purification using standard DNA clean up columns (that do not remove long oligos) can be used. However, we observed that inefficient PCR amplification resulting in the presence of significant contamination of the final product with M1 and M2 tagging oligos can potentially lower the yield of integration at the correct loci (data not shown).

#### Cas12a delivery

We also tested if Cas12a could be delivered using mRNA or protein instead of plasmid-borne Cas12a expression. We found that transformation required electroporation and that for all three expression systems successful tagging could be achieved (Fig. S3c). This indicates the modularity of the system but for the sake of simplicity we used plasmid-borne expression for the remainder of the study.

#### Length of homology arms

From yeast it is known that approx. 28 to 36 nucleotides (nt) of continuous sequence homology are minimally required for homologous recombination of transfected DNA with the genome (Rothstein, 1991). For PCR tagging in yeast, homology arms between 45 and 55 nt in length are routinely used. To obtain some insights into the requirement in mammalian cells, we tested the integration efficiency as a function of the length of the homology arms. This revealed that already short homology arms of 30 nt on both sides allow efficient integration of the cassette (Fig. 3a), but increasing the length results in more efficient integration.

**Figure 3.**
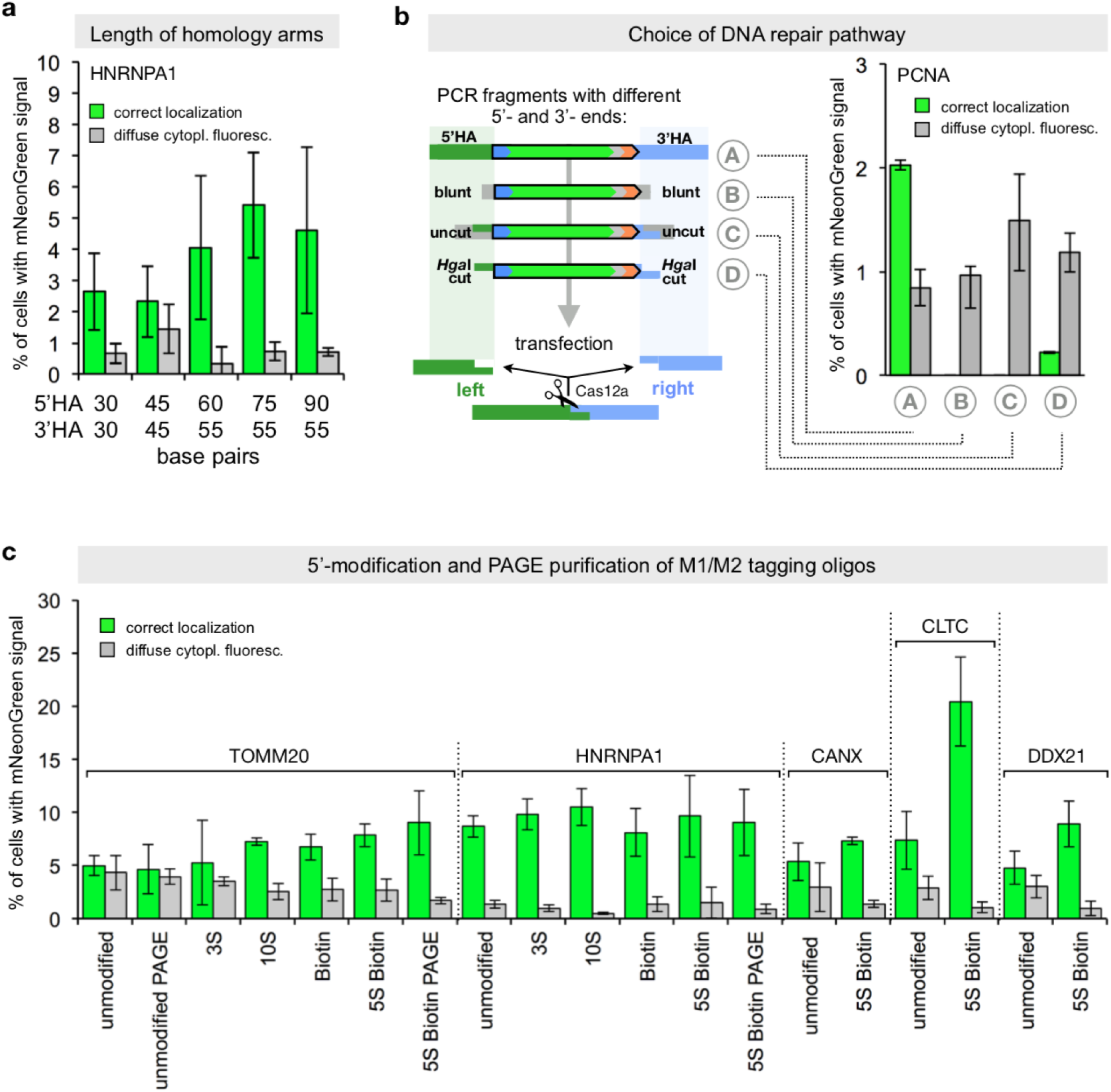
Tagging efficiency as a function of different parameters. (**a**) Length of homology arms. M1 and M2 tagging oligos containing the indicated sequence lengths of homology arm (5’-HA and 3’-HA, respectively) to the destination locus were used for PCR tagging of the HNRNPA1 locus in HEK293T cells. Tagging efficiency was estimated 3 days after transfection as described before. Data from three replicates is shown. Error bars indicate SD. (**b**) PCR cassettes containing various types of ends to direct the choice of DNA repair pathway: Homology arms (90-bp and 55-bp homology, for HR; *A*), blunt ended arms without homology to the target locus (blunt; *B*), *Hga*I cut (*D*) and uncut ends (*C*). Cutting with the type IIS restriction enzyme *Hga*I results in 5-nt 3’-overhangs that are complementary to the overhangs generated by the crRNA directed Cas12a-cleavage of the destination locus. Tagging efficiency was estimated three days later as described in (a) using HEK293T cells. Data from three replicates is shown. Error bars indicate SD. (**c**) Use of modified and/or purified oligos. M1/M2 tagging oligos with the indicated number of phosphorothioate bonds and/or biotin as indicated were used for generation of PCR cassettes. All oligos were ‘cartridge’ purified except for the ones denoted with ‘PAGE’, which were size selected using polyacrylamide gel electrophoresis. Tagging efficiency was estimated three days after transfection as described before using HEK293T cells. Data from three replicates is shown. Error bars indicate SD.

#### Dependence on homology arms

Our control experiment (Fig. 1e) suggested that PCR tagging depends on the presence of homology arms. However, it could still be that a fraction of the productive events is not mediated by HR, but by alternative DNA repair pathways. To test this directly we generated a series of PCR cassettes with different types of ends. In particular, we also generated a PCR cassette with compatible overhangs for direct ligation, by using a Type IIS restriction enzyme (*Hga*I). This enzyme generates ends that contain 3’ overhangs of 5 nt on both sides, which were designed such that they are compatible with the ends produced by Cas12a (Zetsche et al., 2015) in the corresponding genomic locus (Fig. 3b). We observed in-frame integration of the *Hga*I cut fragment, but with lower frequency when compared to the integration in the presence of homology arms (Fig. 3b). This demonstrates the requirement of homology arms for efficient integration. Insertion of the PCR cassettes via c-NHEJ can be observed, but it is rather inefficient.

#### Modified oligonucleotides

End-to-end joining of transfected dsDNA inside cells can be reduced when bulky modifications such as biotin are introduced at the 5’-end of the DNA fragment. This has been reported to enhance targeting efficiency ∼2-fold in *Medaka* (Gutierrez-Triana et al., 2018) and the biotin-modification could contribute to enhance targeting efficiency in mouse embryos (Gu et al., 2018), leading to the insertion of preferentially one copy of the donor DNA. We tested M1/M2 tagging oligos with multiple phosphorothioate bonds (to prevent exonuclease degradation) with and without biotin at the 5’-end. Synthetic oligo synthesis occurs in the 3’ to 5’ direction, and oligo-preparations without size selection are contaminated by shorter species without the 5’-modifications. Therefore, we additionally included size selected (PAGE purified) oligos.

In all cases we observed a 2-to 3-fold reduced frequency of cells with diffuse cytoplasmic fluorescence (Fig. 3c). This is consistent with the idea that the modifications are partially effective in suppressing end-to-end ligation and therefore concatemer formation. Quantification of the targeting efficiency revealed for TOMM20, CLTC and DDX21 an increased tagging efficiency to a maximum of 2-to 3-fold. It was irrelevant, whether the oligos were size-selected or not. However, for HNRNPA1 and also CANX the modifications only slightly enhanced tagging efficiency.

Taken together, these experiments demonstrate the robustness of the procedure and dependency on homology arms for efficient recombination with the target locus, leading to the tagged gene. The use of modified oligos with phosphorothioate exhibits an overall positive effect on tagging efficiency and reduces diffuse cytosolic fluorescence most likely by reducing end-to-end ligation of fragments by c-NHEJ.

### Tagging fidelity and off-target integrations

Integration of DNA by homologous recombination in the genome of mammalian cells might be associated with mutations caused by the integration process or that result from faulty oligos (Fig. 4a). In addition, integration by c-NHEJ and off-target integration of the cassette elsewhere in the genome might occur. To investigate this in more details, we transfected HEK293 cells with PCR cassettes targeting three different genes. The Cas12a cleavage sites were selected to have different positions around the stop codon, either before (CANX), after (HNRNPA1) or directly at the STOP codon (CLTC) (Fig. 4b). For all cases we used protected primers (5S biotin, Fig. 3c) to reduce the load of concatemerized cassettes. Insertion junctions at the targeted gene were amplified from unselected cell populations 18 days after transfection.

**Figure 4.**
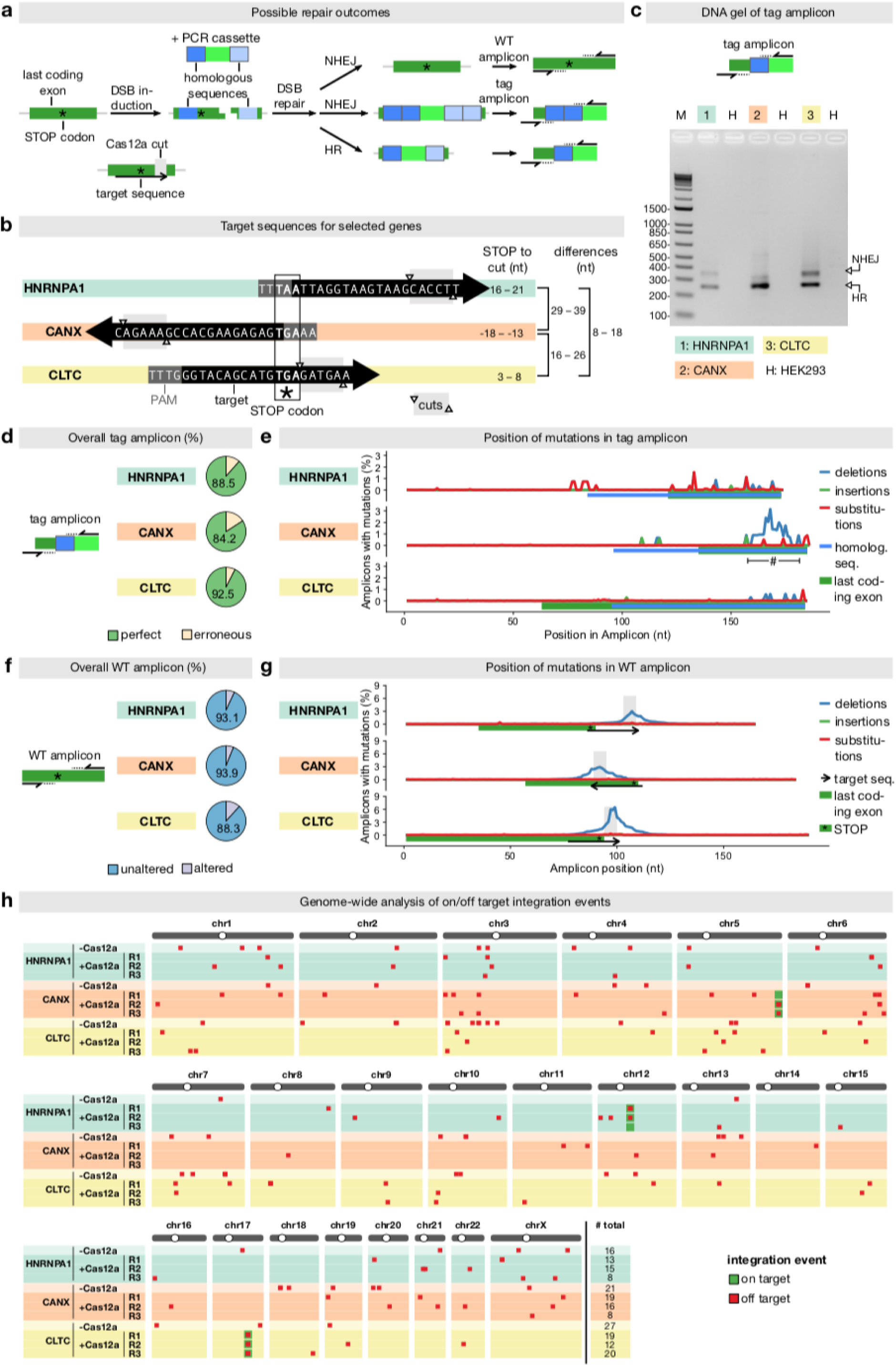
Fidelity of tag integration and off-target events in unselected cell populations. (**a**) Schematic of representation possible repair outcomes following a Cas12a cut at the target site: Cassette integration by HR, integration by c-NHEJ, and DSB repair without cassette integration. (**b**) Target sequences for three selected genes and the resulting distances between induced DSB and the stop codon of the gene. (**c**) PCR amplification of the insertion junction of the respective genes (tag amplicon). HEK293 cells were transfected with mNeonGreen containing PCR cassettes and PCR was performed six days post-transfection. The upper band corresponds to junctions generated by insertion via c-NHEJ and the lower via HR as indicated. (**d**) Sequencing of tag amplicons formed by HR of the same genes as in (c) (>10,000 reads per gene), but using cells 18 days after transfection. The frequencies of reads exhibiting perfect and erroneous exon-tag junctions are given. (**e**) The position of observed mutations in the tag amplicons. The approx. 1.5-2 fold higher frequency of mutations in the insertion junction observed for CANX is caused mostly by small deletions and can be explained by reconstitution of a crRNA targeting site after tag integration with the non-canonical PAM site ‘CCTG’ in the CANX-mNeonGreen fusion. (**f**) Amplification of the crRNA cleavage site of unmodified alleles in cells of (d). The frequencies of reads exhibiting unaltered and altered sequences when compared to the wild type sequence are given. (**g**) Samples as in (f). The position and frequency of specific types of mutations across all reads is shown. (**h**) Off-target integration events detected by Anchor-Seq for the selected genes in three biological replicates in the presence of Cas12a and in one biological replicate without Cas12a. Anchor-Seq samples were prepared using cells 30 days post-transfection from HEK293 cells transfected with mNeonGreen-containing PCR cassettes for the indicated genes. # total, number of detected integration sites.

We used PCR to amplify the insertion junction between the 3’ of the ORF and the inserted tag. This yielded two distinct amplicon population. The shorter bands correspond in their size to the junctions formed by HR tag, and longer and less abundant bands corresponding to the size expected from fragment insertions by c-NHEJ (Fig. 4a-c). Despite that PCR of not fully identical fragments can differ in efficiency, the results suggest the more insertion junction formed by HR are present in the population. Illumina dye sequencing of the shorter bands revealed > 80% correct sequences (Fig. 4d) and most other reads contained mutations that were enriched at the end of the homology region near the junction of the tag (Fig. 4e). This suggest that they result from faulty synthesis of the long oligos and that homologous recombination does select against PCR cassettes containing faulty sequences in the region of the homology arms. Similar observations that select against faulty oligos have been made for yeast (Buchmuller et al., 2019), where it is known that mismatch repair systems prevent recombination between short imperfect sequences (Anand et al., 2017).

We next generated amplicons of the wild type loci to quantify the alterations resulting from DSBs which were not repaired by HR with the PCR cassette. Illumina dye sequencing revealed that between 7-12% amplicons contained small deletions close to the positions of the Cas12a induced DSBs (Fig. 4f). Depending on the exact position of the Cas12a DSB with regard to the stop codon (Fig. 4b) and the exact manner through which the DSB is repaired via the c-NHEJ machinery, this may cause a modification of the C-terminus of the protein due to a frame shift or altered transcript stability (e.g. due to nonsense-mediated decay) (Fig. 4g).

Next, we used Anchor-Seq to determine potential off-target integrations. We used transfected cells that were passaged for 30 days to minimize PCR cassette derived concatemers. We observed multiple off-target integration events throughout the genome (Fig. 4h). Comparison of the integration sites between replicates and controls without Cas12a plasmid did not identify integration sites that are common between the samples (with the exception of integrations at the target locus). This indicates that the majority, if not all, off-target integration events were caused by random integration of the donor template and not due to off-target activity of Cas12a.

Together, this data indicates that a large fraction of the on-target integration-events yield as a result the expected gene-fusions.

### Selection of clones using antibiotics resistance markers and multi-loci tagging

Next, we generated template plasmids that additionally incorporated selection markers for different antibiotics and used them to generate PCR cassettes for tagging 12 genes, including five genes that we did not tag before (Table S1). PCR cassettes were *Dpn*I or *Fsp*EI to selectively digest the DAM methylated template plasmid DNA (which also contains the selection marker). Selection using either Zeocin or Puromycin resistance yielded cell populations highly enriched in cells exhibiting the correct localization of the fluorescent fusion protein (Fig. 5a). The selected populations still contained cells with the diffuse cytoplasmic fluorescence, but the fraction remained either constant or decreased, consistent with the idea that the transcripts leading to this fluorescence originate predominantly from extrachromosomal concatemers.

**Figure 5.**
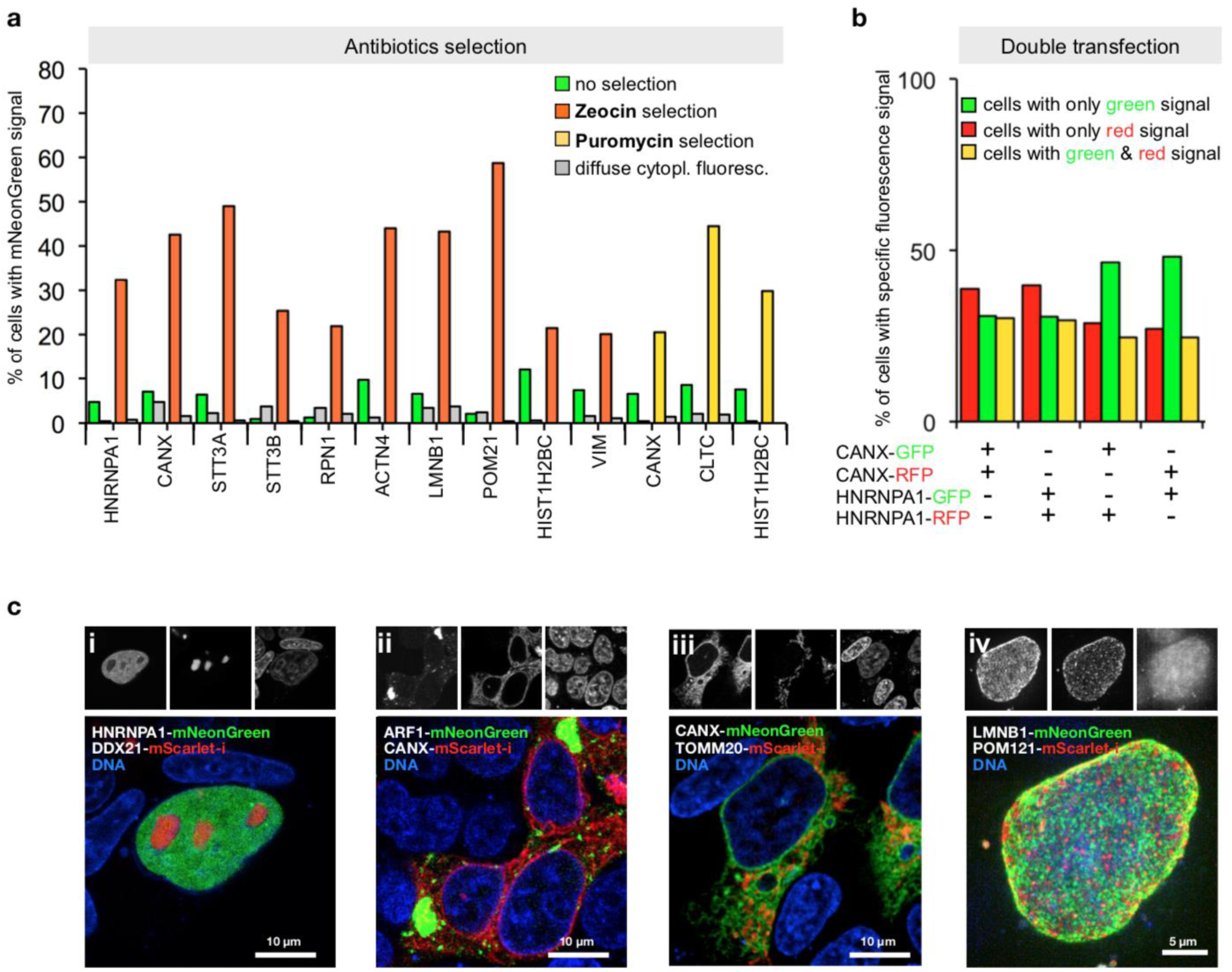
Antibiotic selection and simultaneous tagging of two loci. (**a**) Enrichment of HEK293T cells expressing correctly localized fusion proteins using Zeocin or Puromycin selection as indicated. Antibiotics selection was started three days after transfection. Fractions of cells exhibiting localized or diffuse cytoplasmic fluorescence are shown. Data from one representative experiment is shown. (**b**) Double transfection of cells using PCR cassette reporters for the indicated genes and with the indicated fluorescent protein. For counting, only cells exhibiting correctly localized fluorescence signals were considered (ER localization for CANX tagging, nuclear localization for HNRNPA1 tagging, see Fig. S5). Data from one representative experiment is shown. (**c**) Double tagging of the genes indicated in the images. Representative cells are shown. (i to iii) single plane images, (iv) a maximum projection of multiple planes spanning the upper half of a cell nucleus is shown.

After enrichment of positive cells by Zeocin selection, we isolated individual clones for detailed analysis. PCR identified in all clones correct insertion junctions on the side of the fluorescent protein tag, and in four out of five also on the other side of the PCR cassette. Antibodies detected the corresponding mNeonGreen fusion protein (Fig. S4). HEK293T cells are aneuploid and appear to have up to five copies of the CANX gene (Lin et al., 2014). We also detected the wild type protein of CANX in all clones, indicating that not all copies were tagged (Fig. S4). We used PCR to investigate the presence of concatemers and found that this was the case for four of the clones. Therefore, it appears that correctly tagged clones contain frequently integrated concatemers at the tagged locus, as also predicted from previous work (Folger et al., 1985; 1982). Clones with concatemer might be enriched during antibiotics selection due to the presence of multiple resistance genes. In either case, the inserted additional copies are unlikely to interfere with the tagged gene since they are insulated from the inserted tag by a proper transcription terminator.

To gain insight into the frequency of multiple tagging events, we next generated for CANX and HNRNPA1 two PCR cassettes each, one for tagging with the red fluorescent protein mScarlet-I (Bindels et al., 2017) and one with mNeonGreen, respectively. The resulting four cassettes were then co-transfected into HEK293T cells in mixtures of pairs of two, using all four possible red-green and gene-gene combinations. This detected three types of cells, with green, red, or green and red fluorescence in the nucleus or the ER respectively, as shown for the example of the *HNRNPA1-mScarlet-i/HNRNPA1-mNeonGreen* transfection (Fig. S5). The frequency of each of the three types of cells was roughly equal, no matter whether the same or two different genes were tagged (Fig. 5b). This indicates high double tagging efficiency of different loci, and demonstrates that often more than one allele is tagged. This suggests applications of PCR tagging for the analysis of protein-protein interactions using epitope tagging, or protein co-localization using different fluorescent proteins. We validated this possibility in double tagging experiments (Fig. 5c), which demonstrated simultaneous detection of various cellular structures with one transfection.

Together, this analysis demonstrates that all positive clones contain insertions by HR that yield the correct fusion protein. Insertions are not necessarily single copy, but likely concatenated segments of PCR cassettes. Nevertheless, since the PCR cassette provides STOP codon and a 3’-UTR along with the tag, the generated transcript is properly defined.

### Applications of PCR tagging in different cell lines

So far, we have described and characterized mammalian PCR tagging as a robust workflow for chromosomal tagging in HEK293T and HEK293 cells. To challenge the general applicability of PCR tagging, we tested additional human but also murine cell lines to target genes already tagged successfully in our initial experiments. In each cell line we identified for most genes cells that showed correctly localized green fluorescence. However, we note that for some of these cell lines transfection efficiency was in the lower range so that we observed a tagging frequency of 0.2 to 5% (Fig. 6a-d). Examples of tagged murine myoblast (C2C12) cells are shown in Fig. S6a. For HeLa cells, that also provide only moderate transfection levels, we additionally subjected the cells to selection, and found up to 40% of cells exhibiting the correct localization (Fig. S6b). In conclusion, these results demonstrate that PCR tagging works for different mammalian cell lines and species, including differentiated and stem stem cells, whereby combining transfection with selection vastly increases tagging efficiency.

**Figure 6.**
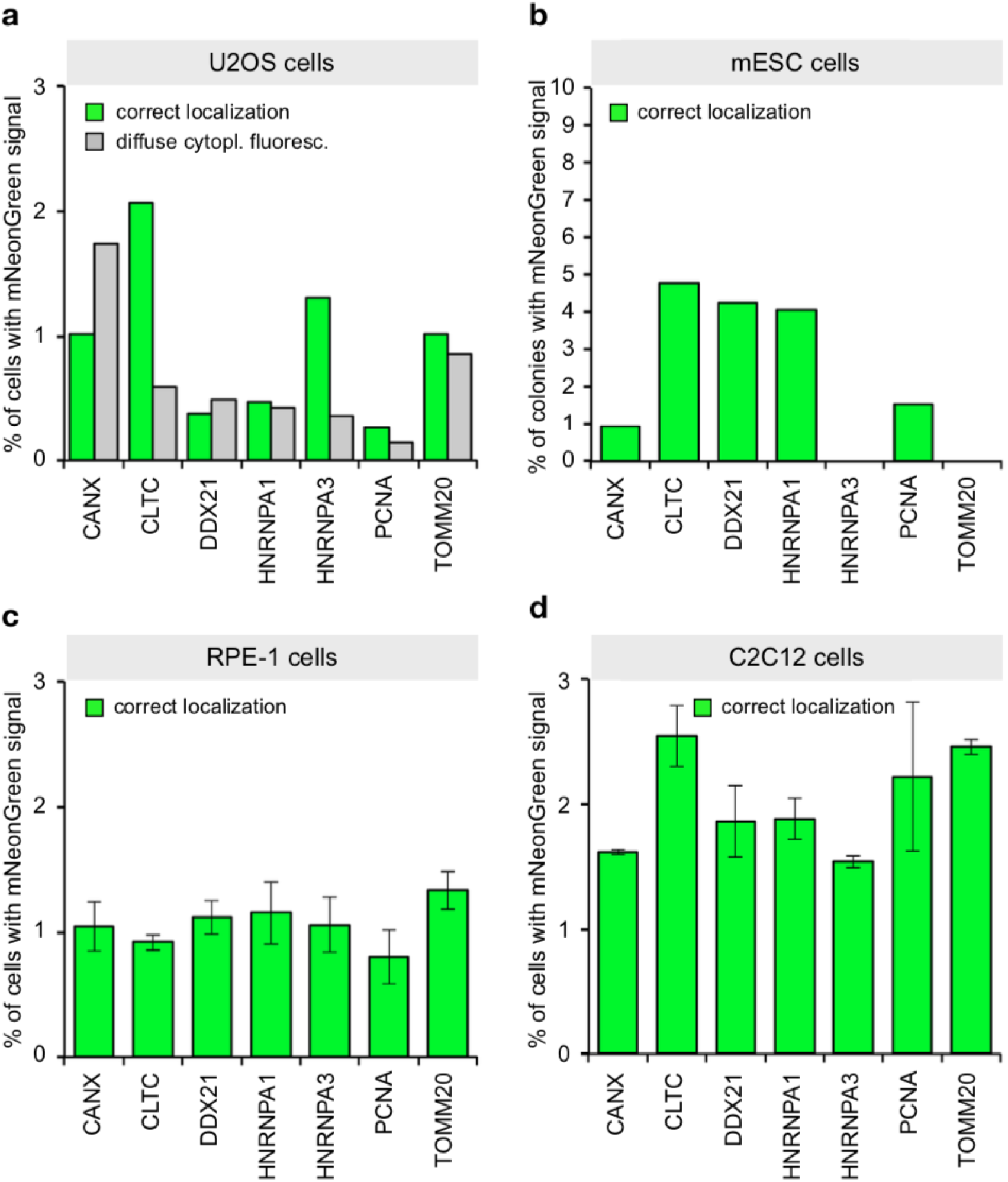
PCR tagging in different cell lines. (**a**) Transfection of U2OS cells using Lipofectamine 2000. After three days the cells were analyzed using HOECHST staining and live cell imaging. Data from one representative experiment is shown. (**b**) Electroporation of mESC cells with PCR cassettes for tagging the indicated genes. After three days the cells were fixed using paraformaldehyde and analyzed. We counted microcolonies that have at least one positive cell. Please note: For these cells we did not quantify cells with diffuse cytoplasmic fluorescence, since paraformaldehyde fixation prior to imaging leads to an increase in cellular background fluorescence. This prevented the detection of the weak cytoplasmic diffuse mNeonGreen fluorescence. Data from one representative experiment is shown. (**c**) Electroporation of RPE-1 cells. Cells were analyzed two days later. Experimental setup similar to (b). Data from three replicates is shown. Error bars indicate SD. (**d**) Electroporation of C2C12 cells. Cells were analyzed two days later. Experimental setup similar to (b). Data from two replicates is shown. Error bars indicate SD.

### crRNA design, PAM site selection and genomic coverage

Next, we asked how well Cas12a-targeted PCR tagging covers the human genome. Our tagging approach relies on relatively short homology arms of the PCR cassette. This constrains the target sequence space, since cleavage of the target locus must be inside the area of the homology arms, leaving enough sequence for recombination. In addition, insertion of the cassette needs to destroy the crRNA cleavage site, in order to prevent re-cleavage of the locus (see also legend to Fig. 4g). For C-terminal protein tagging these criteria confine potentially useful protospacer-associated motif (PAM) sites to a region of 17 nt on both sides of the STOP codon including the STOP codon, with the PAM site or protospacer sequence overlapping the STOP codon (Fig. 7a). So far, we have used Cas12a from *Lachnospiraceae bacterium* ND2006 (LbCas12a) (Zetsche et al., 2015), but PAM sites that are recognized by this Cas12a (TTTV) (Gao et al., 2017) and that are located in this area of a gene are relatively infrequent and would allow C-terminal tagging of about one third of all human genes (Fig. 7b). To increase this number, we first tested different Cas12a variants with altered PAM specificities (Gao et al., 2017). The results demonstrated that other variants and PAM sites are also functional and can be used for PCR tagging (Fig. 7c). Considering these and additional enCas12a variants (Gao et al., 2017; Kim et al., 2016a; Tóth et al., 2018; Kleinstiver et al., 2019; Sanson et al., 2019) renders approx. 97% of all human genes accessible for C-terminal PCR tagging (Fig. 7b). To increase the number of suitable PAM sites for one Cas12a variant further we extended the search space into the 3’-UTR (typically 50 nt) (Fig. 7a) and adjusted the design of the M2 tagging oligo such that a small deletion occurs that removes the binding site of the crRNA. Since tagging introduces a generic terminator for proper termination of the tagged gene, this small deletion is unlikely to have an impact on the tagged gene. Considering the extended search space and the currently available palette of Cas12a variants we calculated that close to 100% of all human ORFs (Fig. 7b) are amenable for C-terminal PCR tagging.

**Figure 7.**
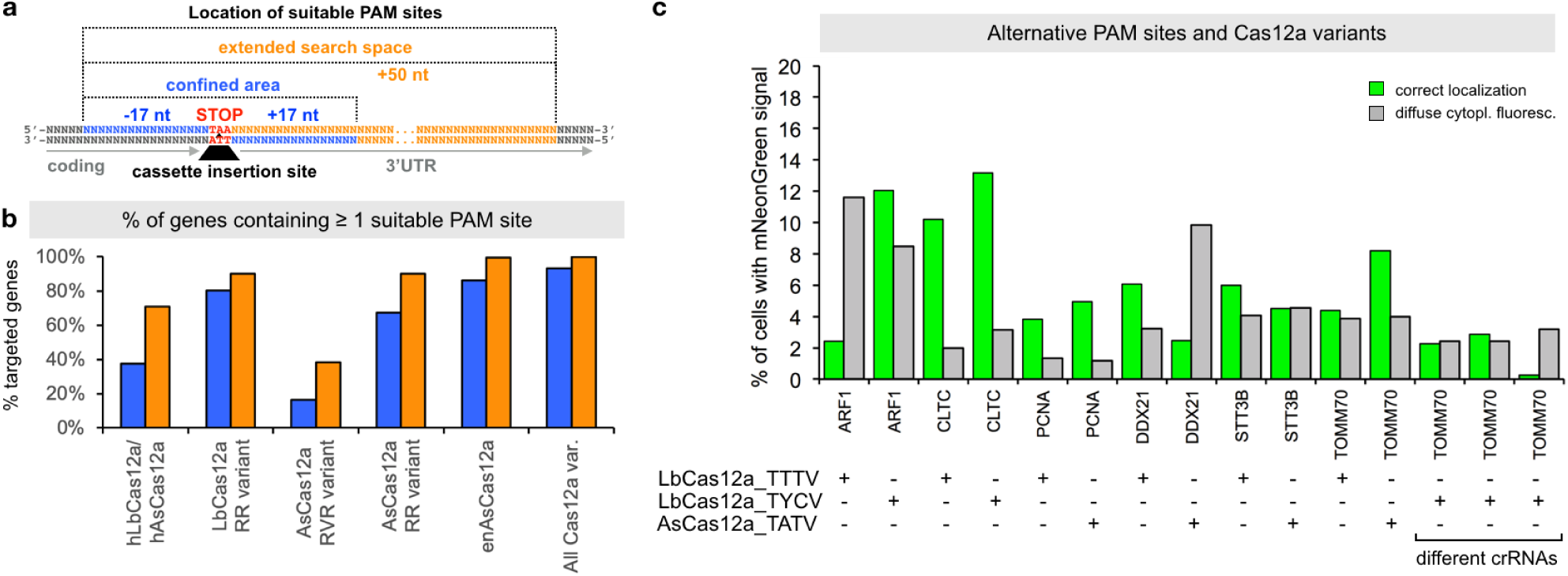
PCR tagging enables C-terminal tagging of the majority of human genes. (**a**) Search space for Cas12a-PAM sites suitable for C-terminal protein tagging. PCR cassette insertion into the genome using PAM sites located in the confined search space (blue) lead to a disruption of the crRNA target sequence. This would not be the case for PAM sites in the extended search space (orange). To prevent re-cleavage after insertion the homology arm of the PCR fragment (provided by the M2 tagging oligo) is designed such that a small deletion in the region after the STOP codon does lead to the disruption of the crRNA target site. (**b**) Fraction (in %) of human genes with suitable PAM sites near the STOP codon, as a function of the confined and extended search spaces (a) and different Cas12a variants as indicated. For calculation, we used the following PAM sites: hLbCas12a/hAsCas12a: TTTV; LbCas12a RR variant: TYCV, TYTV; AsCas12a, RVR variant: TATV; AsCas12a RR variant: TTTV, TYCV; enAsCas12a: TTYN, VTTV, TRTV, VTCC, HSCC, TACA, TTAC, CACC (Tier 1 and 2 PAM sites). (**c**) Tagging of the indicated genes in HEK293T cells. Helper plasmids with different Cas12a genes, as indicated. PCR cassettes contained crRNA genes with matching PAM site specificity. For *TOMM70* three different Cas12a variants were tested, using three different crRNA sequences for AsCas12a, as indicated. Tagging efficiency was determined three days after transfection. Data from one representative experiment is shown.

### PCR tagging toolkit for mammalian cells

To further facilitate application of mammalian PCR tagging, e.g. for quick C-terminal fluorescent protein labeling, we set up a webpage for oligo design (Fig. 1a). The online tool (www.pcr-tagging.com) requires as input the ‘Ensembl transcript ID’ (www.ensembl.org) of the target gene. Alternatively, the genomic DNA sequence around the desired insertion site, i.e. the STOP codon of the gene of interest for C-terminal tagging can be provided. The software then generates the sequence of the M1 tagging oligo, which specifies the junction between the gene and the tag. Next, the software identifies all PAM sites for the available Cas12a variants and uses these to generate crRNA sequences and to assemble corresponding M2 tagging oligos. M2 tagging oligos are designed such that the integration of the PCR cassette does lead to a disruption of the crRNA binding site or PAM site in order to prevent re-cleavage of the locus. M2 tagging oligos are then ranked based on the quality of the PAM site and the presence of motifs that might interfere with crRNA synthesis or function. M1/M2 tagging oligos can be used with template plasmids based on different backbones: either without a marker, with Zeocin or Puromycin resistance genes (Fig 8a). We generated a series of template plasmids containing different state of the art reporter genes (Table 1, examples shown in Fig. 8b).

**Table 1.** Overview of the available template plasmids for PCR tagging. For additional information, see Supplementary Table 2.

| Tag | Type | Oligos | Markerless | Zeocin | Puromycin | Application |
| --- | --- | --- | --- | --- | --- | --- |
| moxBFP | fluorescent protein | M1/M2 | pMaCTag-01 | pMaCTag-Z01 | pMaCTag-P01 | Imaging |
| TagBFP | fluorescent protein | M1/M2 | pMaCTag-02 | pMaCTag-Z02 | pMaCTag-P02 | Imaging |
| moxCerulean3 | fluorescent protein | M1/M2 | pMaCTag-03 | pMaCTag-Z03 | pMaCTag-P03 | Imaging |
| mTFP1 | fluorescent protein | M1/M2 | pMaCTag-04 | pMaCTag-Z04 | pMaCTag-P04 | Imaging |
| eGFP | fluorescent protein | M1/M2 | pMaCTag-05 | pMaCTag-Z05 | pMaCTag-P05 | Imaging |
| sfGFP | fluorescent protein | M1/M2 | pMaCTag-06 | pMaCTag-Z06 | pMaCTag-P06 | Imaging |
| mNeonGreen | fluorescent protein | M1/M2 | pMaCTag-07 | pMaCTag-Z07 | pMaCTag-P07 | Imaging |
| mClover3 | fluorescent protein | M1/M2 | pMaCTag-08 | pMaCTag-Z08 | pMaCTag-P08 | Imaging |
| mVenus | fluorescent protein | M1/M2 | pMaCTag-09 | pMaCTag-Z09 | pMaCTag-P09 | Imaging |
| mKOk | fluorescent protein | M1/M2 | pMaCTag-10 | pMaCTag-Z10 | pMaCTag-P10 | Imaging |
| mScarlet-i | fluorescent protein | M1/M2 | pMaCTag-11 | pMaCTag-Z11 | pMaCTag-P11 | Imaging |
| mScarlet | fluorescent protein | M1/M2 | pMaCTag-12 | pMaCTag-Z12 | pMaCTag-P12 | Imaging |
| mCherry | fluorescent protein | M1/M2 | pMaCTag-13 | pMaCTag-Z13 | pMaCTag-P13 | Imaging |
| mNeptune2.5 | fluorescent protein | M1/M2 | pMaCTag-14 | pMaCTag-Z14 | pMaCTag-P14 | Imaging |
| miRFP670 | fluorescent protein | M1/M2 | pMaCTag-15 | pMaCTag-Z15 | pMaCTag-P15 | Imaging |
| Dronpa3 | fluorescent protein | M1/M2 | pMaCTag-16 | pMaCTag-Z16 | pMaCTag-P16 | Imaging |
| mEos4b | fluorescent protein | M1/M2 | pMaCTag-17 | pMaCTag-Z17 | pMaCTag-P17 | Imaging |
| mKeima8.5 | fluorescent protein | M1/M2 | pMaCTag-18 | pMaCTag-Z18 | pMaCTag-P18 | Imaging |
| mCherry-mNeonGreen | fluorescent protein | M1/M2 | pMaCTag-19 | pMaCTag-Z19 | pMaCTag-P19 | Imaging |
| 10xSunTag | detection tag | M1/M2 | pMaCTag-20 | pMaCTag-Z20 | pMaCTag-P20 | Imaging |
| 24xSunTag | detection tag | M1/M2 | pMaCTag-21 | pMaCTag-Z21 | pMaCTag-P21 | Imaging |
| SNAP | chem. modification | M1/M2 | pMaCTag-22 | pMaCTag-Z22 | pMaCTag-P22 | Imaging |
| Halo | chem. modification | M1/M2 | pMaCTag-23 | pMaCTag-Z23 | pMaCTag-P23 | Imaging |
| TAP | Affinity tag | M1/M2 | pMaCTag-24 | pMaCTag-Z24 | pMaCTag-P24 | PPIs |
| YFP-LID | degron | M1/M2 | pMaCTag-25 | pMaCTag-Z25 | pMaCTag-P25 | function |
| mAID-Clover | degron | M1/M2 | pMaCTag-26 | pMaCTag-Z26 | pMaCTag-P26 | function |
| 1xHA | antibody epitope | M1/M2 | pMaCTag-27 | pMaCTag-Z27 | pMaCTag-P27 | detection |
| 3xHA | antibody epitope | M1/M2 | pMaCTag-28 | pMaCTag-Z28 | pMaCTag-P28 | detection |
| Strep-II | Affinity-tag | M1/M2 | pMaCTag-29 | pMaCTag-Z29 | pMaCTag-P29 | detection |
| VSV-G | antibody epitope | M1/M2 | pMaCTag-30 | pMaCTag-Z30 | pMaCTag-P30 | detection |
| V5 | antibody epitope | M1/M2 | pMaCTag-31 | pMaCTag-Z31 | pMaCTag-P31 | detection |
| S-Tag | bimolecular compl. | M1/M2 | pMaCTag-32 | pMaCTag-Z32 | pMaCTag-P32 | detection |
| FLAG | antibody epitope | M1/M2 | pMaCTag-33 | pMaCTag-Z33 | pMaCTag-P33 | detection |
| 3xFLAG | antibody epitope | M1/M2 | pMaCTag-34 | pMaCTag-Z34 | pMaCTag-P34 | detection |

**Figure 8.**
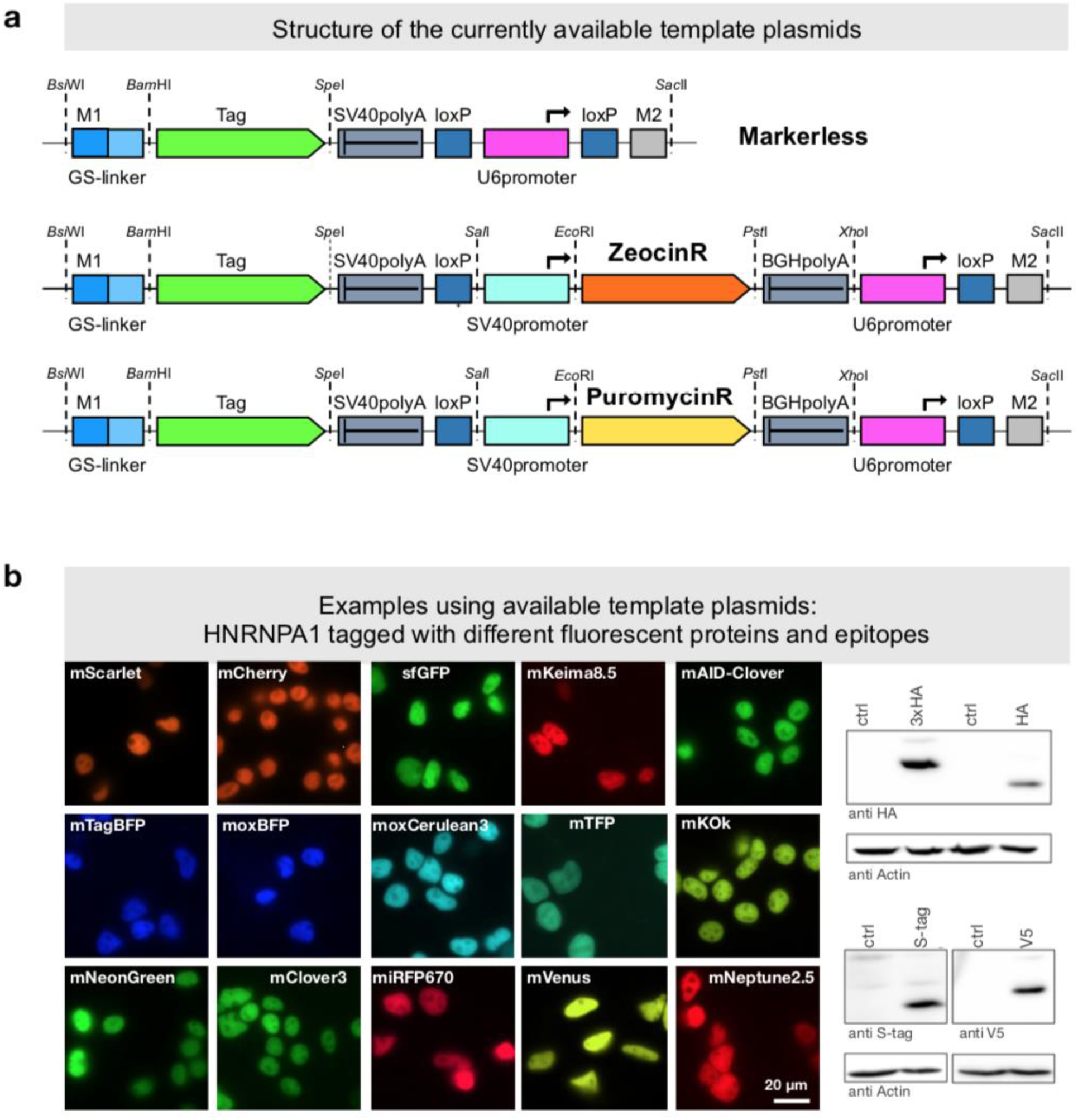
PCR tagging Toolkit for mammalian cells. (**a**) Schematic outline of the template plasmids provided. (**b**) Examples of HNRNPA1 tagging using different available cassettes. Complete list of features and sequence files are provided in Table 1, S2. Western blot analysis was performed three days after transfection with crude lysate of a cell pool. Fluorescence microscopy was performed using cells three days after transfection.

Ongoing efforts continue to improve optimal crRNA prediction and to eliminate crRNAs with potential off-target binding activity. The current version of the server already allows to flexibly add novel Cas12a variants, by adjusting PAM site specificity and the sequence of the corresponding constant region of the crRNA.

In conclusion, Cas12a-mediated PCR tagging of mammalian genes using short homology arms is a rapid, robust and flexible method enabling endogenous gene tagging. The versatility of the method suggests many types of applications for functional or analytical gene and protein studies in mammalian cells.

## Discussion

In this paper we demonstrate efficient targeted integration of DNA fragments of several kbp in size into the genome of mammalian cells, guided by short homology arms (<100 bp). Integration is assisted by CRISPR-Cas12a and a crRNA that is expressed from the DNA fragment itself. This enables a PCR-only strategy for the production of the gene specific reagents for tagging. In addition, we present a software tool for oligo design and established streamlined procedures for application in several cell lines.

PCR tagging can be easily scaled up and parallelized – since it needs only two oligos per gene. In yeast, where PCR tagging is very efficient even in the absence of targeted DSB induction, the ease of upscaling permitted the creation of many types of genome-wide resources. In these all genes were modified in the same manner, i.e. by gene deletion or by tagging with a fluorescent protein or affinity tag (Gavin et al., 2002; Ghaemmaghami et al., 2003; Huh et al., 2003; Meurer et al., 2018; Winzeler et al., 1999). We believe that in mammalian cell culture similar endeavors are now at reach using the approach presented in this study.

Tagging efficiency might be influenced by various factors including chromatin structure and expression levels. Our choice of relatively highly expressed genes as convenient reporters to validate and investigate the method might bias the efficiency. While genome wide analysis of tagging efficiency for Cas12a tagging using yeast did not reveal a correlation of expression levels and tagging efficiency (Buchmuller et al., 2019), further experiments will be needed to validate whether this is also the case in mammalian cells.

The use of tagged genes always raises the question about the functionality of the tag-fusion. Here, two questions matter: How does tagging affect gene regulation, and how does it affect protein function? Many aspects of protein tagging have been discussed in literature, i.e. from functional or structural points of view. But ultimately, one has to be aware of the fact that a cell expressing a tagged gene is a mutant, and that the tag does not necessarily correctly report about the behavior of the untagged protein (Lundberg and Borner, 2019). As part of good laboratory practice this demands for some sort of phenotypic analyses to investigate the functionality of the tagged gene/protein and/or orthogonal experiments to obtain independent validation of the conclusions that were derived with the tagged clone(s). In haploid yeasts, genome wide analysis of the influence of a C-terminal tag revealed that >95% of the ∼1000 essential yeast genes, when endogenously tagged with a large tag such as a fluorescence protein reporter, retain enough functionality to not cause an obvious growth phenotype under standard growth conditions (Khmelinskii et al., 2014).

Various methods for gene tagging with long DNA fragments in mammalian cells have been developed, including methods that are tailored for particular DNA damage pathways such as c-NHEJ or HR to repair induced DSBs via CRISPR-Cas9 or other endonucleases. In all cases the heterologous sequence to be inserted needs to be provided either by using circular or linear repair templates generated *ex vivo* or *in vivo* upon endonuclease excision of the repair template (Agudelo et al., 2017; He et al., 2016; Lackner et al., 2015; Merkle et al., 2015; Suzuki et al., 2016; Zhang et al., 2017; Zhu et al., 2015; Roberts et al., 2017; Chen et al., 2018). In non-germline cells, the insertion precision by HR is often limited due to the co-existence of alternative repair pathways, and errors such as small indels are frequently observed near one or the other side of the inserted fragment. Therefore, a substantial number of clones need to be screened in order to obtain a few correct ones (Koch et al., 2018). PCR tagging does not generate seamlessly integrated tags, since it is accompanied by a generic transcription termination site that replaces the endogenous 3’-UTR. This actually bears the advantage that it reduces the errors associated with tag insertion, since an erroneous insertion downstream of the PCR cassette, i.e. caused by c-NHEJ instead of homologous recombination will only affect the 3’-UTR of the gene, which is not used for the tagged allele. Obviously, this constitutes a compromise, and includes the possibility that important gene regulatory sequences are omitted from the tagged gene, e.g. miRNA binding sites, targeting motifs or sequences regulating mRNA stability. While for mammalian cells no global data set about the regulatory impact of the 3’-UTR on gene expression is available, data from yeast, where seamless tagging was compared with tagging using a generic 3’-UTR, demonstrated that only about 11% of the genes were impacted in their expression more than 2-fold (Meurer et al., 2018).

Based on our detailed analysis of three genes in Fig 4c and Fig 5a we conclude that c-NHEJ is not the dominating repair outcome after all and that it is easy to obtain enriched populations containing the correct gene fusions. Given the fact that enriched populations are composed from many different clones, it is possible to use them for a rapid assessment of experimental questions, for example the localization of one or the co-localization of even two proteins upon endogenous expression, in a specific condition, environment or cell line, by simply scoring multiple cells. Since they are derived from different clones, clone-specific effects can be spotted rapidly and considered in the analysis. This avoids the need of perfectly characterized cell lines with exactly the intended genomic modification and does save a lot of time.

DSB induction at off-target locations by Cas12a *in vivo* has been investigated extensively in the past and found to be reduced compared to canonical Cas9 variants (Kleinstiver et al., 2016; Kim et al., 2016b; 2017). In agreement with this, our unbiased targeted NGS experiment (Fig. 4h) detected no obvious off-target activity related to Cas12a. Nevertheless, we observed random integrations of the PCR cassettes which is common when foreign DNA is introduced into mammalian cells (Folger et al., 1982; Saito et al., 2017). Analysis of multiple independent clonal cell lines will exclude unwanted effects from off-target integrations.

The toolset available for PCR tagging can easily be expanded by constructing new template plasmids. Maintaining a certain level of standardization such as the preservation of the primer annealing sites for the M1 and M2 tagging oligos in new template cassettes, makes it is possible to reuse already purchased M1/M2 tagging oligos of the same gene for many different tagging experiments. We recommend the use of chemically modified M1 and M2 primers (e.g. with 5S and biotin) as we noticed considerable enhancement in tagging efficiency.

Further improvements of the tagging efficiency might be possible, i.e. by targeting the repair template to the CRISPR endonuclease cut site (Roy et al., 2018), or by using Cas12a variants that are only active in S/G2 phase of the cell cycle (Smirnikhina et al., 2018).

Beyond mammalian cells, there may be other species where this strategy could improve tagging methodology, i.e. many fungal species that require a DNA double strand break for targeted integration of a foreign DNA fragment.

In conclusion, PCR-mediated C-terminal gene tagging is a simple, non-commercial, easily adoptable method to exploratively study protein localization or to explore other functional aspects using endogenous-level expression. It is simple to design the oligos (www.pcr-tagging.com) and open access to all other resources is granted (via Addgene or colleagues), and reagents can be freely exchanged. We believe that for many applications PCR tagging is quicker than the construction of a plasmid for transient transfection or exogenous chromosomal integration.

With PCR tagging at hand, many different and exciting experimental avenues are becoming possible, from the rapid assessment of protein localizations to high throughput localization studies of many proteins.

## Supporting information

Supplementary Table S1 - list of tagged genes

Supplementary Table S2 - detailed template plasmid info

Supplementary Table S3 - Plasmids

Supplementary Table S4 - Oligos

## Author contribution

M.K. and M.M. designed the project and together with M.K.L, J.F. and K.H. designed the experiments. J.F., K.H., M.M., B.K., J.D.K. and D.K. performed the experiments. K.H. analyzed the NGS data, K.G. wrote the web-tool for primer design. All authors analyzed the data and discussed the results. M.K. wrote the manuscript with input from all authors.

## Acknowledgements

The authors wish to thank Cyril Mongis for help with IT infrastructure, Anne Schlaitz and Frauke Melchior for critical reading of the manuscript. We acknowledge support by the Deutsche Forschungsgemeinschaft (DFG, grant KN498/12-1), the Collaborative Research Center SFB1036, the state of Baden-Württemberg through bwHPC for high-performance computing and SDS@hd for data storage (grant INST 35/1314-1 FUGG). K.H. was supported by a HBIGS graduate school fellowship and J.D.K. by a fellowship of the Boehringer Ingelheim Fonds. G.P. and B.K are supported by the Collaborative Research Center SFB873 and the Heisenberg Program of the DFG (granted to G.P.).

## Materials and Methods

### Plasmids and oligos

Plasmids are listed in Table 1, S2 and S3. Sequences are provided for download from www.knoplab.de. pMaCTag Plasmids can be obtained from www.addgene.org. All used oligos for cloning, Anchor-Seq and gene tagging are listed in Table S4.

### Construction of template cassettes

For cloning standard restriction enzyme digests or oligo annealing and ligations using enzymes from NEB was used. Most of the elements inside the template cassettes (M1-mNeonGreen-SV40polyA-ZeocinR-BGHpolyA-hU6promoter) were custom synthesized (gBlock, IDT) and cloned via *Bsi*WI and *Xba*I into a *Bsi*WI and *Spe*I cut pFA6a backbone. The SV40 promoter was cloned separately into the cassette via *Sal*I and *Eco*RI, since it contains repeats and could not be synthesized together with the other elements. In addition to the ZeocinR marker we have also introduced a PuromycinR marker. Because the standard DNA sequence for this marker is very GC-rich and difficult to amplify by PCR, we synthesized a new version with lower GC-content and cloned it via *Eco*RI and *Pst*I into the cassette. To obtain a cassette without a marker the SV40promoter-ZeocinR-BGHpolyA sequence was removed using a digest with *Sal*I and *Xho*I and subsequent religation of the backbone. This resulted in three different plasmids based upon the backbone pFA6 (see Fig. 8a).

The mNeonGreen ORF of these template plasmids is flanked by unique restriction sites and is therefore easily exchangeable. For introduction of new tags *Bam*HI and *Spe*I sites can be used. For a high flexibility in cloning, the sticky ends of both restriction sites are compatible to sticky ends produced by other enzymes (*Bcl*I/*Bgl*II and *Avr*II/*Nhe*I/*Xba*I, respectively).

All tags listed in Table 1 and Table S2 are cloned either by amplification from template plasmids with oligos containing restriction sites or by annealing of two oligos and are ligated into *Bam*HI/*Spe*I cut backbones of pMaM523/526/541 (for detailed information see Table S2) to retrieve template cassettes called pMaCTag (**p**lasmid for **Ma**mmalian **C**-terminal **Tag**ging) with the following naming scheme:

pMaCTag-xy: Tag xy, no marker, pMaM526 backbone

pMaCTag-**Z**xy: Tag xy, **ZeocinR** marker, pMaM523 backbone

pMaCTag-**P**xy: Tag xy, **PuromycinR** marker, pMaM541 backbone

### M1 and M2 tagging oligo design

The online oligo design tool (www.pcr-tagging.com) was implemented using Shiny. The interactive web application was developed in R v3.4.4 (R Core Team, 2014) with the R packages shiny v1.1.0 (Chang et al., 2018) and shinyjs v1.0 (Attali, 2017). The R package Biostrings v2.46.0 (Pagès et al., 2018) is used for searching PAM sites. The latest code is available from our GitHub repository (www.github.com/knoplab). Oligo design principles are as follows:

#### M1 tagging oligo

The design of the M1 tagging oligo is straight forward as it contains only two functional elements: the primer annealing site for PCR, which is constant in all template cassettes (TCAGGTGGAGGAGGTAGTG), and the sequence of the homology arm, which is derived from the target locus.

#### *Example*: M1 tagging oligo (for TOMM70)

##### Description of elements

**5’-homology (90 bases before the insertion site, direct orientation)---primer annealing site for PCR**

##### Sequence

ATGGAGATGGCCCATCTGTATTCACTTTGCGATGCCGCCCATGCCCAGACAGAAGTTGCAAAGAAATACGGATTAAAACCACCAACATTATCAGGTGGAGGAGGTAGTG

#### M2 tagging oligo

The design of the M2 tagging oligo is more complex. It contains the annealing site for PCR (GCTAGCTGCATCGGTACC), the direct repeat sequence of the crRNA, which is Cas12a-variant specific, and the protospacer sequence of the crRNA, which depends on available PAM sites at the target locus, a terminator for the Pol III RNA polymerase and the homology arm, as outlined below.

#### *Example*: M2 tagging oligo (for TOMM70)

##### Description of elements

**3’-homology (55 bases after the insertion site, reverse orientation)---Pol III terminator---crRNA protospacer sequence---<u>crRNA direct repeat sequence</u>---primer annealing site for PCR**

##### Sequence

CAGTTGAAGAGGGGGTAAACTTTTAAAAAGAGGGTCAGTCTGCTTTCCCCCTGTTAAAAAAAG TCTGCTTTCCCCCTGTTT<u>ATCTACAAGAGTAGAAATTA</u>GCTAGCTGCATCGGTACC

Criteria used for ranking crRNAs currently implemented in www.pcr-tagging.com, listed according to priority:

1. Location of the crRNA binding site in the genome in a region where it becomes destroyed upon cassette integration in order to prevent re-cleavage. This can be on either side in close proximity of the insertion site (17 nt up and downstream of the insertion site). If no suitable crRNA binding site is found in this confined search space, the software offers the option to select PAM sites in the 3’-region of the insertion site (extended search space). In this case the design of the homology arm of the M2 tagging oligo is adjusted in such a manner that the target site of the crRNA is deleted. This results in a small deletion in the 3’-UTR of the gene after the insertion site of the cassette. Since the PCR cassette contains a transcriptional terminator, we deem this to be non-critical. With these criteria, it is possible to design suitable crRNAs for C-terminal tagging of the vast majority of mammalian genes (Fig. 7b).
2. The protospacer sequence should preferably not contain four or more ‘T’-s in a row, since this might lead to premature termination of the Pol III transcription of the crRNA (Arimbasseri et al., 2013) In practice, we observed that crRNAs with ‘TTTT’ are frequently functional.
3. PAM sites are ranked according to literature (Gao et al., 2017; Kim et al., 2016a; Tóth et al., 2018; Kleinstiver et al., 2019; Sanson et al., 2019). In addition, unconventional PAM sites were considered (MCCC for the AsCas12a RR variant and RATR for LbCas12a RVR variant), based on depositor comments on the Addgene webpage. For ranking crRNAs, conventional PAM sites are preferred.
4. If multiple crRNAs are fulfilling these criteria, they are ranked according to the position of the cleavage site, with a preference for greater distance after the STOP codon.

### Synthesis of M1 and M2 tagging oligo

All M1 and M2 tagging oligos were obtained from Sigma-Aldrich using a 0.05 µmole synthesis scale and are RP1 cartridge purified, unless otherwise stated (as in Fig. 3c).

### PCR of template cassettes using M1 and M2 tagging oligos

PCR using long oligos is not always easy and requires optimized protocols. We routinely use a self-purified DNA polymerase for PCR (Pfu-Sso7d (Wang et al., 2004)). Alternatively, for cassette PCR also commercial high-fidelity polymerases can be used. We have tested Phusion (ThermoFisher) and Velocity polymerase (Bioline). We note that the Phusion polymerase using the buffer provided by the manufacturer does not work for PCR cassette amplification with M1 and M2 tagging oligos, whereas good amounts can be obtained using our buffer. Velocity polymerase works using the buffer provided by the manufacturer.

We found that all polymerases work well using the buffer conditions and amplification scheme shown below, yielding similar amounts of PCR cassette.

PCR mixture:

- 5.0 µL of 10x HiFi-buffer (200 mM Tris-HCl, pH 8.8; 100 mM (NH_4_)_2_SO_4_, 500 mM KCl, 1% (v/v) Triton X-100, 1 mg/mL BSA, 20 mM MgCl_2_)
- 5.0 µL of dNTPs (10 mM stock, Bioline, BIO-39026)
- µL of MgCl_2_ (50 mM stock)
- 5.0 µL of betaine (5 M stock, Sigma-Aldrich, 61962)
- 0.3 µL of template DNA (200 ng/µL stock)
- 2.5 µL of M1 tagging oligo (10 µM stock)
- 2.5 µL of M2 tagging oligo (10 µM stock)
- x µL of H_2_O up to 50 µL
- 1 µL self-purified DNA polymerase (1 U/µL) or 0.5 µL Phusion or 0.25 µL Velocity polymerase PCR was mixed on ice and was carried out in a Biometra TRIO (Analytik Jena) using the following program:
- 3 min at 95 °C
- 30 cycles of:

- 20 s at 95 °C
- 30 s at 64 °C
- XX s at 72 °C (45 s per kb) (see Table S2)
- 5 min at 72 °C
- 4 °C
- After PCR, 0.4 µL *Dpn*I or *FspEI* (and 1.67 µL Enzyme activator) was added to the reaction mixture and incubated at 37 °C for 1 h to digest the template that contains a selection marker that would contaminate the transfection.
- PCR products were analyzed by agarose gel electrophoresis and purified using column purification (Macherey-Nagel).

#### Note

Sometimes a particular pair of oligos does not yield a product upon PCR. In this case it is worth testing whether adding 2 min. on top of the calculated elongation time does solve the problem. If not, it might be that synthesis of the primer went wrong. To determine the faulty primer, pair-wise PCR with established M1 and M2 primers can be used to identify the faulty primer. Usually, ordering the same primer again solves the problem. Providers may wave the cost of re-ordering.

### Preparation of genomic DNA

Genomic DNA for Experiments shown in all Figures except Fig. 4 was isolated from HEK293T cells using a protocol adapted from Ref. (Greene and Sambrook, 2012). After washing with PBS, a confluently grown 6-well was lysed in 600 µL SNET buffer (20 mM Tris pH 8.0, 400 mM NaCl, 5 mM EDTA pH 8.0, 1% SDS) and 2 µL of RNase A (10 mg/mL RNAse A, 10 mM Tris-HCl pH 8.0, 10 mM MgCl_2_) was added for 30 minutes at room temperature. Afterwards Proteinase K (20 mg/mL Proteinase K, 50 mM Tris-HCl pH 8.0, 1.5 mM CaCl_2_, 50% glycerol) was added for another 30 minutes at room temperature. Proteins were precipitated using 200 µL 3 M K-Acetate solution, followed by precipitation of the DNA with Isopropanol and washing with 70% Ethanol. DNA was dried and dissolved in TE (10 mM Tris, 1mM EDTA) buffer. Genomic DNA for experiments shown in Fig. 4 were purified according to the instructions of the manufacturer using the High Pure PCR Template Preparation Kit (Roche) followed by RNAse A digest and a final purification with the High Pure PCR Product Purification Kit (Roche).

### Targeted next-generation sequencing of tagged and wild type alleles

Tag-and wild type-specific amplicons from cells used in the experiments shown in Fig. 4 were generated from 200 ng genomic DNA (gDNA) using junction-specific primers (Table S4) by a two-step nested PCR with Velocity polymerase (Bioline). The first PCR reaction was performed for 15 cycles with 60 °C annealing and 30 s elongation and then purified with AMPure XP PCR beads (Beckman Coulter). The second PCR was performed for 15 cycles for wild type-specific and for 21 cycles for tag-specific amplicons respectively using 60°C annealing and 30 s elongation. PCR products were size-selected by gel electrophoresis on 2% Agarose/TAE and gel extracted by column purification (Macherey-Nagel). Amplicons were paired-end sequenced with 500 cycles on a MiSeq system (Illumina) using the Amplicon-EZ (150-500 bp) service by Genewiz to acquire at minimally 13,123 reads per sample. Paired-reads were merged and aligned to the respective expected amplicon references using CRISPResso (v2.0.29) (Kleinstiver et al., 2019) with parameters “cleavage_offset”: 1 and “window_around_sgrna”: 0. Mutations were subsequently quantified using a custom R script excluding primer binding sites in the analysis.

### Next-generation sequencing of genomic DNA with Anchor-Seq

Sequencing libraries for experiments to determine cassette junction sites presented in Fig. 2 were prepared based on our previously published Anchor-Seq protocol (Meurer et al., 2018) with some modifications to the adapter design to include unique molecular identifiers (UMIs) (Table S4) (Buchmuller et al., 2019). Quantified libraries were sequenced paired-end with 300 cycles on a NextSeq 550 sequencing system (Illumina) with a spike-in of 20% phiX gDNA library (Illumina). Raw reads were trimmed from technical sequences (adapter and cassette sequences) using custom scripts (Julia v0.6.0 and BioSequences v0.8.0). The trimmed reads were aligned to the human reference genome (Genome Reference Consortium Human Build 38 for alignment pipelines, ftp://ftp.ncbi.nlm.nih.gov/genomes/all/GCA/000/001/405/GCA_000001405.15_GRCh38/seqs_for_alignment_pipelines.ucsc_ids/) using bowtie2 (v2.3.3.1) (Langmead and Salzberg, 2012). Template cassette sequences were included in the reference genome as decoy. Aligned reads were grouped with UMI-tools (Smith et al., 2017) based on unique molecular identifiers included in the Anchor-Seq adapters. Enriched integration sites were further evaluated and counted using IGV (v2.4.10) (Robinson et al., 2011).

Sequencing libraries for mapping the genomic integration sites of off-target integrations presented in Fig. 4 were prepared by a modified Anchor-Seq protocol using tagmentation instead of sonication for gDNA fragmentation (Picelli et al., 2014). In detail, 100 ng/µL Tn5(E54K,L372P) transposase (Hennig et al., 2018) was loaded with 2.5 µM annealed adapters (P5-UMI-gri501…506-ME.fw, Tn5hY-Rd2-Wat-SC3) in 100 mM Tris-HCl (pH 7.5) by incubating the reaction for 1h at 23 °C. Tagmentation reactions with 1 µg gDNA were prepared mixed with tagmentation buffer (10 mM Tris-HCl, pH 7.5; 10 mM MgCl_2_, 25% (v/v) DMF). For our batch of Tn5 transposase we achieved reasonable tagmentation using an enzyme:gDNA mass ratio of 0.35. Tagmentation reactions were completely used as input for a first PCR reaction with cassette-and Tn5-adapter-specific primers (5Btn-hmNeong.rv, P5.fw) with NEBNext Q5 HotStart polymerase (New England BioLabs) with 15 cycles of 68 °C and 1 min elongation. Biotinylated amplicons were enriched using Dynabeads MyOne Streptavidin C1 beads (Invitrogen) according to the manufacturers protocol. These beads were then used as input of a second PCR with cassette-and Tn5-adapter-specific primers (P7-gri701…706-hmNeong.rv, P5.fw) with NEBNext Q5 HotStart polymerase (New England BioLabs) with 25 cycles of 68 °C and 1 min elongation. PCR products were size-selected for 400-550 bp using a 2% agarose/TAE gel and column purification (Macherey-Nagel). Libraries were sequenced as above. Raw reads were trimmed as already mentioned, but aligned to the human reference genome supplemented with PCR cassette sequences with bwa mem (v0.7.17-r1188) (Li, 2013). Mapped insertion sites were summarized by a custom R script and further evaluated and counted using IGV (v2.4.10 (Thorvaldsdóttir et al., 2013)).

### In-Vitro Transcription (IVT) of LbCas12a mRNA

Template for IVT of LbCas12a mRNA was amplified from pY016 with primers (CMV-fw, bGH_polyA_IVT.rv (Table S4) using self-purified DNA polymerase. The PCR reaction was column-purified using the Monarch PCR & DNA Cleanup Kit (New England BioLabs). The IVT reaction including DNAseI digest was performed with the mMESSAGE mMACHINE T7 Transcription Kit (Invitrogen) according to the manufacturers instructions. After quality control by gel-electrophoresis the IVT product was purified by phenol-chloroform extraction and subsequent lithium acetate and isopropanol precipitation and the mRNA was reconstituted in nuclease-free water.

### Cell counting and Fluorescence microscopy

*For Fig. 6b-d -*Cells were grown on coverslips (No. 1.5, Thermo Fischer Scientific), washed once with PBS and fixed with 3% PFA for 10 minutes at 37 °C. After fixation, coverslips were washed 3 times with PBS, incubated in PBS containing 0.1 µg/mL 4 ‘, 6-Diamidin-2-phenylindole (DAPI) for 10 minutes and embedded in Mowiol. Coverslips were coated with 0.1% gelatin Type B (Sigma Aldrich) for culturing C2C12 and 0.2% gelatin Type A (Sigma Aldrich) for C2C12 and mES cells, respectively. Images of RPE-1 and C2C12 cells were acquired as Z stacks using Zeiss Axio Observer Z1 equipped with 40x NA 1.3 PlanNeo oil immersion objective, and AxioCam MRm CCD camera using ZEN software. Images of mESC colonies were acquired as Z stacks using Nikon A1R confocal microscope equipped with Nikon Plan Apo λ 20x NA 0.75 objective, using NIS elements software. Maximum intensity projections of the Z stacks were prepared using Fiji (Schindelin et al., 2012) (Schneider et al., 2012).

For cell counting, random fields of view were inspected in the HOECHST/DAPI channel and all nuclei present in the entire field of view were counted. Cells containing transfected fluorescent protein expressing cassettes were then counted subsequently in the same fields of view using the appropriate illumination wavelengths. In some experiments counting was done using images recorded in the same manner.

For Fig. 5c images were taken with Zeiss LSM 780 confocal microscope using a Plan-APOCHROMAT 63x, 1.40 NA Oil Objective (panels i-iii) or a Leica Spinning DMi8 Spinning Disk microscope with HC PL APO 63x, 1.40 NA Oil Objective (panel iv).

For all other Figs. For live cell imaging, cells were splitted 24h after transfection into 8 well µ-slides (Ibidi). Analyses of transfected cells were performed 3 days after transfection or as described in the figure legends. Cells were stained with Hoechst 33342 (4 µg/mL in PBS, Thermo Fisher Scientific) for 5 minutes and then the medium was changed to FluoroBrite (Thermo Fisher Scientific) supplemented with 10% FBS (Gibco) and 20 mM HEPES-KOH, pH 7.4 (Thermo Fisher Scientific). For counting and imaging different microscopes were used: Nikon Ti-E widefield epifluorescence microscope or a DeltaVison with each 60x oil immersion objectives (1.49 NA, Nikon, 1.40 NA, DeltaVision). Z-stacks of 11 planes with 0.5 µm spacing were recorded with 100 ms exposure time. Single plane images and maximum intensity z-projections are shown. Subcellular localizations were identified and scored visually.

### Western Blotting

Cells were solubilized in SDS sample buffer (50 mM Tris-HCl, pH 6.8; 10 mM EDTA, 5% glycerol, 2% SDS, 0.01% bromophenol blue) containing 5% β-mercaptoethanol. All samples were incubated for 15 min at 65 °C. Denatured and fully-reduced proteins were resolved on Tris-glycine SDS-PAGE followed by western blot analysis using the following antibodies: rat monoclonal anti-HA (11867423001; Roche), mouse monoclonal anti-V5 (V8012; Sigma), anti-S-tag mouse monoclonal antibody (MA1-981; Thermo Fisher), rabbit polyclonal anti mNeonGreen Tag (53061S, Cell Signaling), rabbit anti Calnexin (ab22595; abcam).

### Tissue culture

h-TERT-immortalized Retinal Pigment Epithelial (RPE-1, ATCC, CRL-4000, USA) cells were grown in DMEM/F12 (Sigma Aldrich) supplemented with 10% fetal bovine serum (FBS, Biochrom), 2 mM L-glutamine (Thermo Fisher Scientific) and 0.348% sodium bicarbonate (Sigma Aldrich). Mouse myoblast C2C12 cells (gift from Edgar R. Gomis, iMM, Portugal) were grown in DMEM High Glucose (Sigma Aldrich) supplemented with 20% fetal bovine serum (FBS, Biochrom). Mouse embryonic stem cell line E14 (gift from Frank van der Hoeven, DKFZ, Germany) were grown in Knockout DMEM (Thermo Fisher Scientific) supplemented with 10% ESC qualified FBS (Thermo Fisher Scientific), 2 mM GlutaMax (Thermo Fisher Scientific), 0.1 mM β-mercaptoethanol, 10^3^ units of murine leukemia inhibitory factor (LIF from ESGRO, Millipore). mES cells were grown under feeder-free conditions on 0.2% gelatin Type B coated dishes (Sigma Aldrich).

HEK293T, HeLa and U2OS cells were grown in DMEM High Glucose (Life technologies) supplemented with 10% (vol/vol) fetal bovine serum (Gibco).

All cell lines were grown at 37 °C with 5% CO_2_, and regularly screened for mycoplasma contamination.

Selection was performed using 1 µg/mL Puromycin (Sigma Aldrich) or 500 µg/mL Zeocin (Invitrogen) for HEK293T cells. For HeLa cells 300 µg/mL Zeocin was used.

### Transfection

#### Chemical transfection

Transfection of HEK293T, HeLa and U2OS cells was performed using Lipofectamine 2000 (Invitrogen) according to protocol of the manufacturer and using a 24-well format. If not stated otherwise, 500 ng Cas12a Plasmid and 500 ng of the PCR cassette were used for transfection of one well in a 24-well plate.

#### Electroporation

Plasmids containing Cas12a variants and PCR cassettes were electroporated into RPE-1, C2C12, and mESCs using 2 mm gap cuvettes and NEPA-21 electroporator (Nepa Gene, Japan) according to manufacturer’s instructions. OPTI-MEM (Thermo Fisher Scientific) was used as electroporation buffer.

For electroporation of HEK293T cells the Neon Transfection System (Thermo scientific) was used according to the protocol of the manufacturer using 2 pulses of 20 ms and 1150 V.

### Generation of clonal lines

After Zeocin selection cells were trypsinized from a confluent plate and counted in a Neubauer chamber. Three cells per well were calculated and seeded in a 96-well plate. After 5 d wells were checked for single clones. After another 7-10 days cells were checked for fluorescence and positive clones were transferred to a 24-well plate.

## Supplementary Information

### Terminology

Throughout the manuscript and in Table 1 we use the following terms in a consistent manner in order to denote the different components and processes:

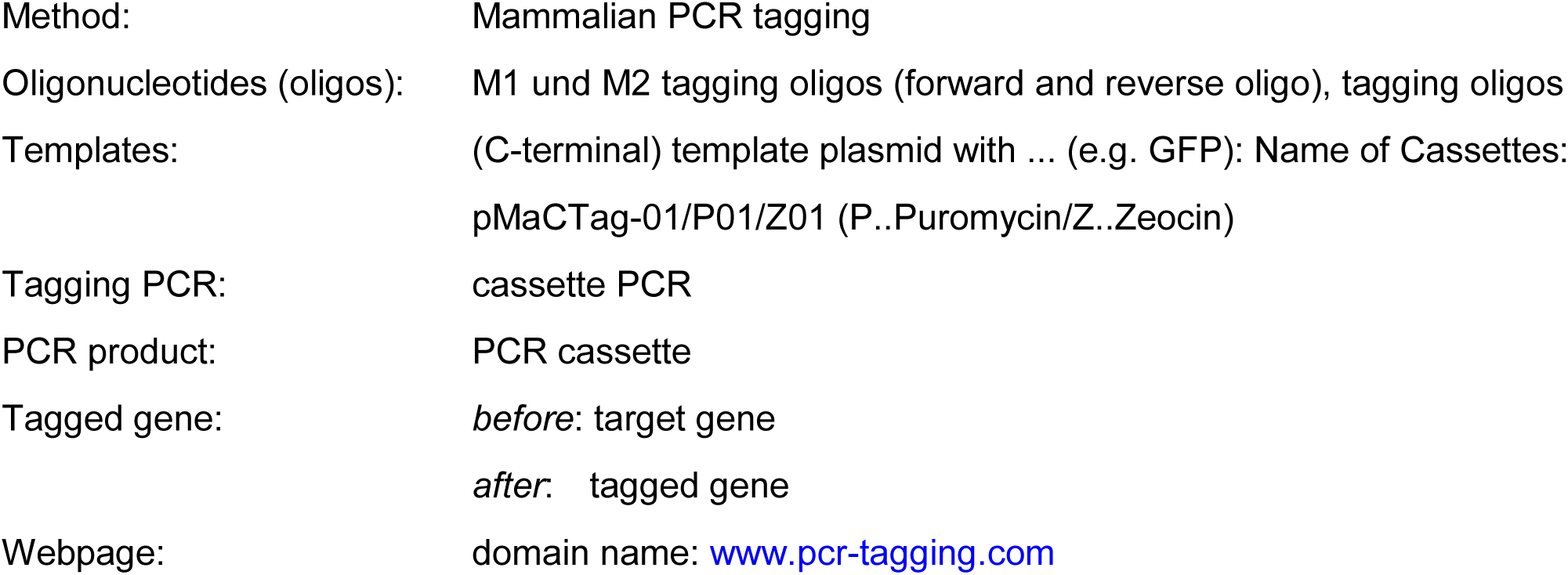

### Supplementary Figures and Tables

**Figure S1.**
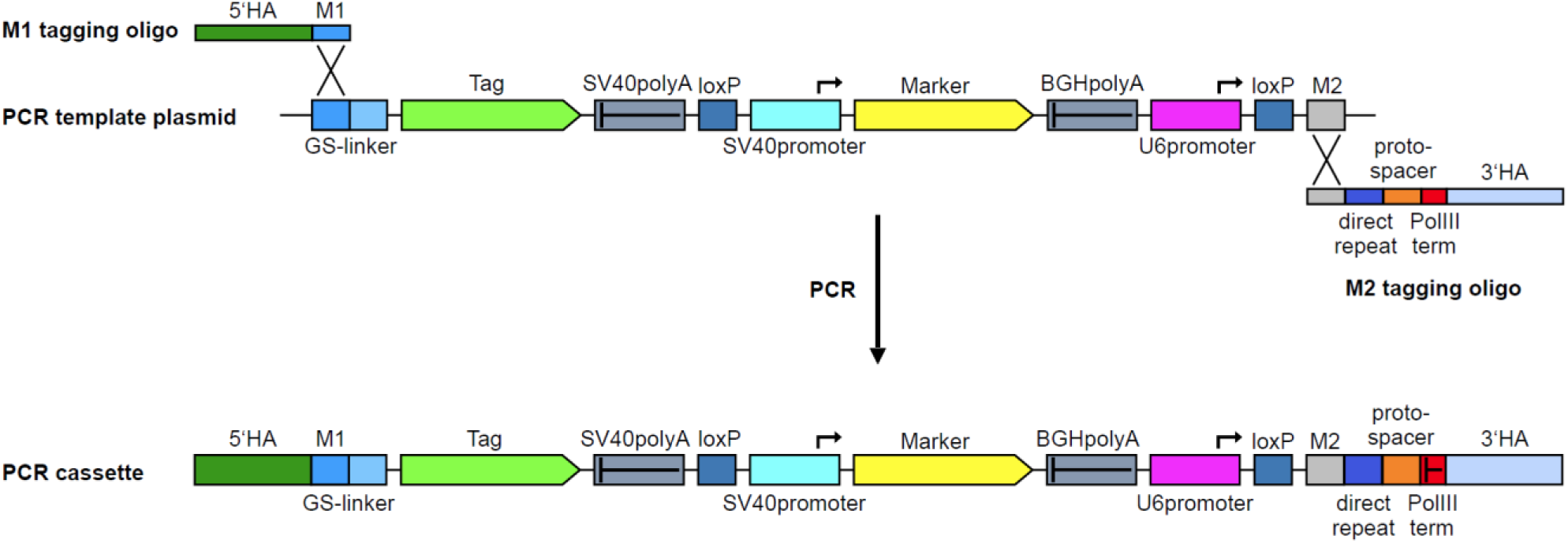
PCR Strategy. PCR is performed with M1 and M2 tagging oligos and a template cassette that contains the tag. The M1 and M2 tagging oligos provide the homology arms (HA, ∼55 to 90 nt in length) for targeted integration. The M2 tagging oligo additionally provides the direct repeat and a protospacer sequence (orange) for a Cas12a endonuclease. The template cassette contains the desired tag and additional features, such as a selection marker. It also contains the U6 Pol III promoter for driving crRNA expression. PCR yields a linear DNA fragment (PCR cassette) that contains homology arms to the target locus and a functional crRNA gene to cleave the locus.

**Figure S2.**
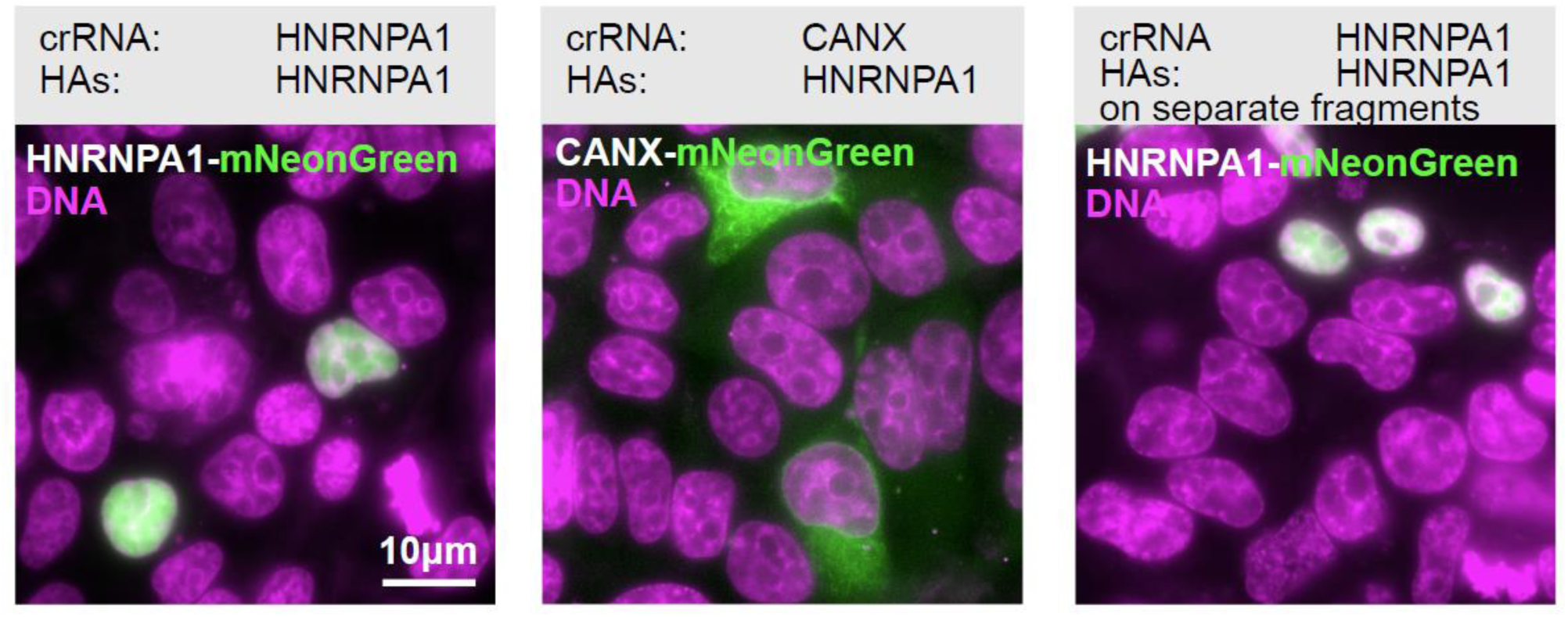
Control transfections (related to Fig. 1f) Control transfections to demonstrate the effect of the crRNA and the presence of homology arms (HAs). Locus specificity of the HAs and the crRNA as indicated.

**Figure S3.**
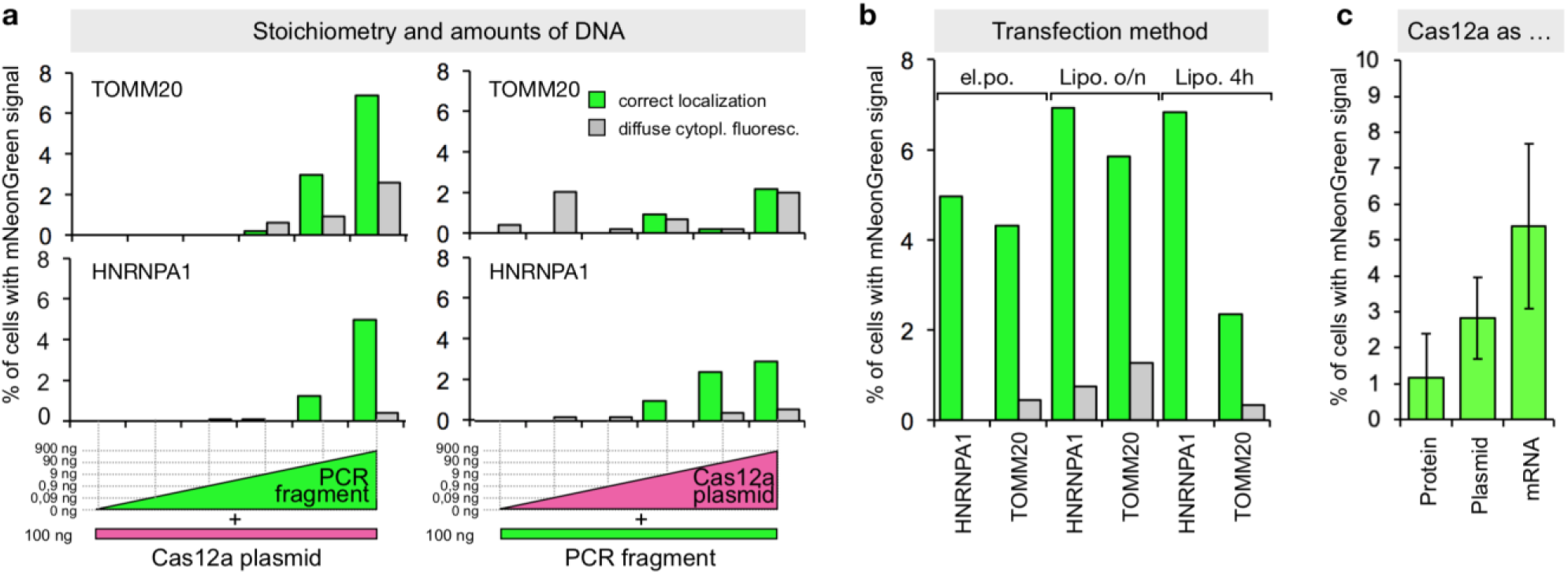
Exploring transfection parameters. (**a**) Impact of transfected amounts of DNA on tagging efficiency using HEK293T cells. Transfected amounts of PCR cassette and Cas12a plasmid as indicated. Always 1 µg of DNA was transfected using lipofectamine. pUC18 was used as neutral DNA. Tagging efficiency was determined three days later by HOECHST staining and live cell imaging. Data from one representative experiment is shown. (**b**) HEK293T cells were transfected for 4 hours or overnight using Lipofectamine 2000 or transfected using electroporation, as indicated. Tagging efficiency was determined three days later as described in (a). Data from one representative experiment is shown. (**c**) HEK293T cells were transfected in duplicates by electroporation with Cas12a protein, Cas12a-encoding mRNA or Cas12-encoding plasmid. For protein-based expression 100 ng of PCR cassette while for mRNA and plasmid 1.5 µg PCR cassette were electroporated. Error bars indicate range between the technical duplicates.

**Figure S4.**
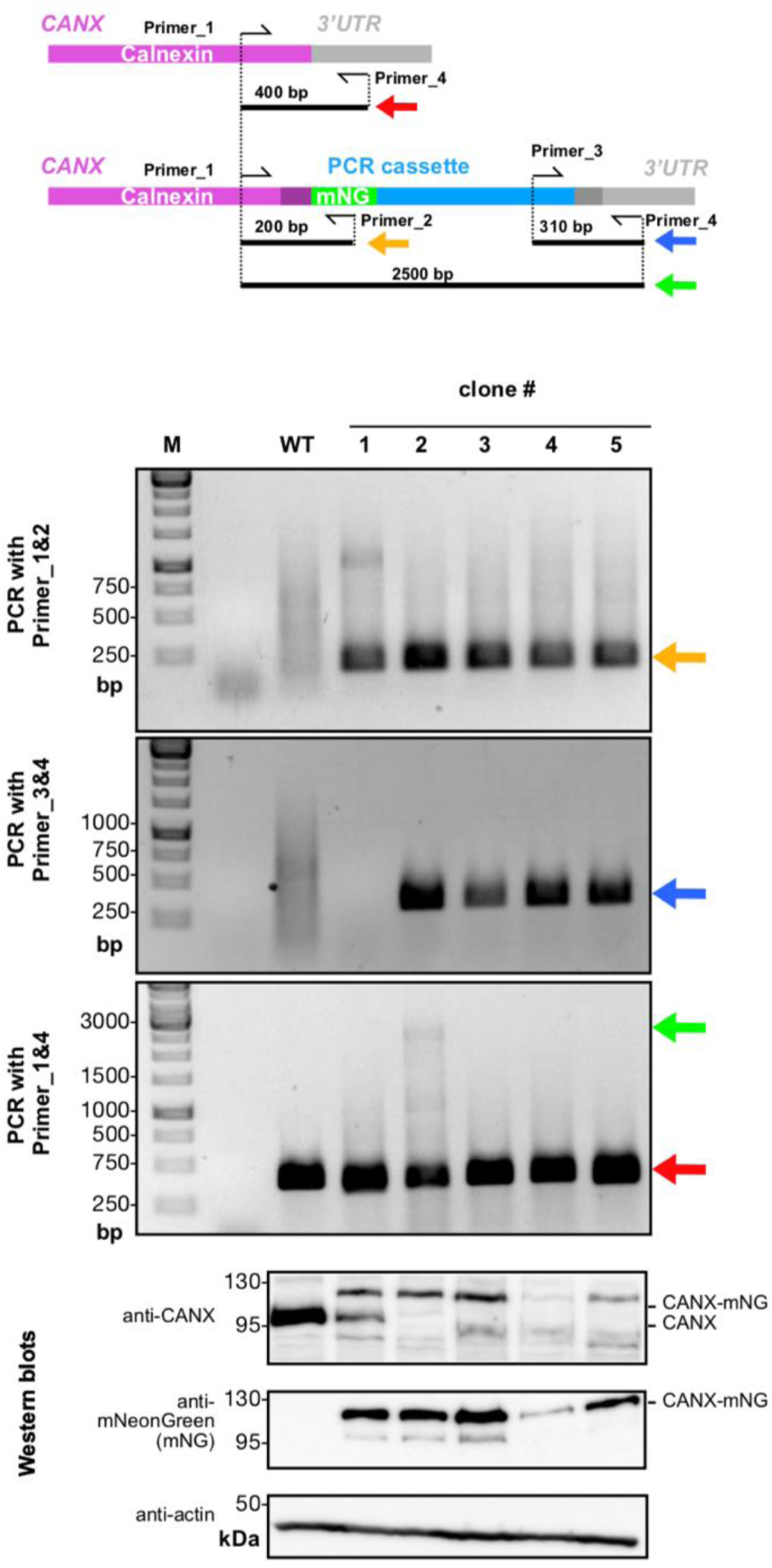
Analysis of clones from a CANX-mNeonGreen tagging experiment. PCR analysis of single clones using primer for PCR of characteristic fragments indicative for correctly inserted fragments. Primer that anneal to chromosomal DNA were chosen to reside outside of the sequences that are contained in the homology arms for recombination. For western blot analysis antibodies specific to mNeonGreen or to Calnexin were used.

**Figure S5.**
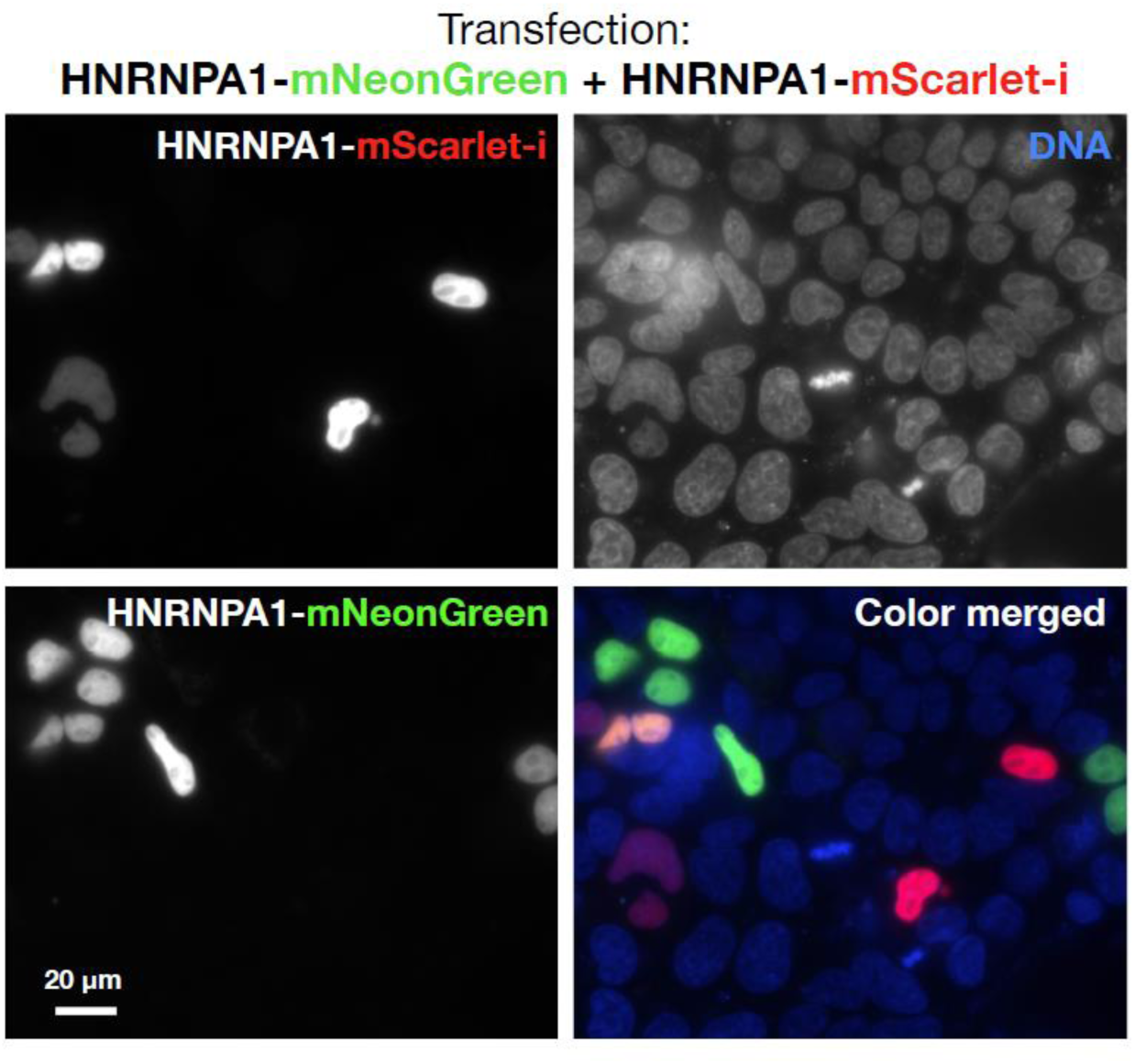
Multi-color integration. Double tagging using a mixture of HNRNPA1-mScarlet-i and HNRNPA1-mNeonGreen PCR cassettes. For analysis, dual color fluorescence images were acquired.

**Figure S6.**
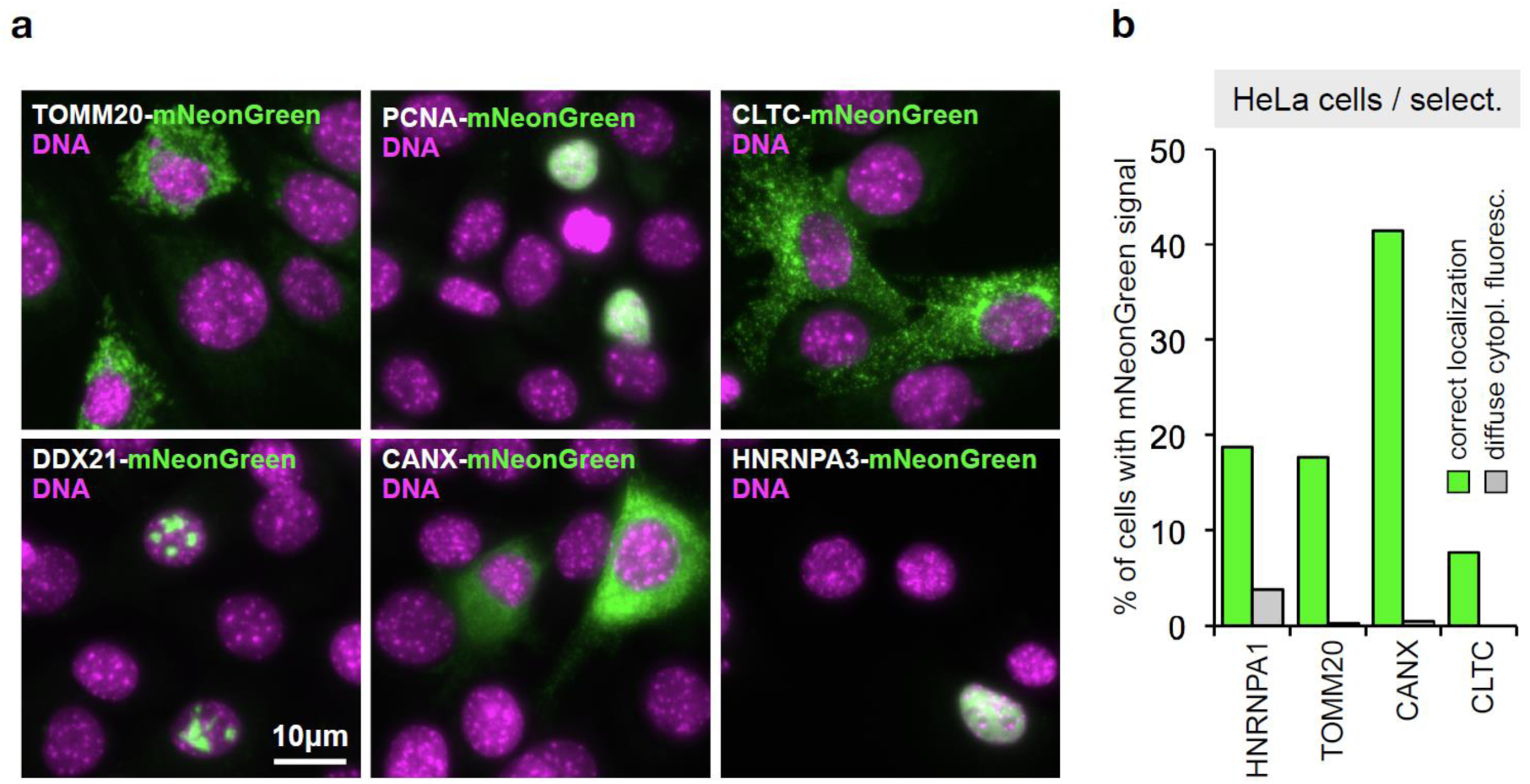
Tagging in C2C12 and Hela cells. (**a**) Sample images from C2C12 cells (Fig. 6d), 5 days after transfection. (**b**) HeLa cells transfected using Lipofectamine 2000. Cells were grown for three days without, and 10 days in the presence of Zeocin using HOECHST staining and live cell imaging. Data from one representative experiment is shown.

