## Supplementary Table S1 - list of tagged genes for "CRISPR-Cas12a-assisted PCR tagging of mammalian genes"

**Supplementary Table S1.** Genes tagged in this study

**Gene	Name	Localization**

CLTC	Clathrin heavy chain	Coated vesicles

CANX	Calnexin	ER

RPN1	Ribophorin 1	ER

RPN2	Ribophorin 2	ER

STT3A	Oligosaccharyltransferase Complex	ER

STT3B	Oligosaccharyltransferase Complex	ER

ARF1	ADP-ribosylation factor 1	Golgi/Endosomes

VDAC1	Voltage-dep. anion-select. channel 1	Mitochondria

TOMM70	Mitochondrial import receptor subu.	Mitochondria

TOMM20	Mitochondrial import receptor subu.	Mitochondria

HNRNPA1	Heterog. nuclear ribonucleoprot. A1	Nucleus

DDX21	Nucleolar RNA helicase 2	Nucleoli

HNRNPA3	Heterog. nuclear ribonucleoprot. A3	Nucleus

SSB	Sjogren  Syndrome  Antigen  B	Nucleus

PCNA	Proliferating cell nuclear antigen	Nucleus

SLC25A17	Peroxisomal memb. prot. PMP34	Peroxisome

LMNB1	Lamin B1	Nuclear inner membrane

ACTN4	α-Actinin-4	Actin cytoskeleton

POM21	POM121 Transmembrane Nucleoporin	Nuclear envelope

HIST1H2BC	Histone Cluster 1 H2B Family Member C	Chromatin

VIM	Vimentin	Intermediate filaments
