## Supplementary Table S3 - Plasmids for "CRISPR-Cas12a-assisted PCR tagging of mammalian genes"

**Name	Backbone	Description	Reference**

pFA6a	-	backbone of template cassettes	{Wach:1994ut}

pY016	pcDNA3.1	pcDNA3.1-hLbCpf1	{Zetsche:2015bp}

pY210	pcDNA3.1	pcDNA3.1-hAsCpf1(TYCV)	{Gao:2017gn}

pY220	pcDNA3.1	pcDNA3.1-hAsCpf1(TATV)	{Gao:2017gn}

pY230	pcDNA3.1	pcDNA3.1-hLbCpf1(TYCV)	{Gao:2017gn}

pMaCTag-Zxy	pFA6a	see Table 1	This study

pMaM518	pFA6a	GS_linker-mNeonGreen-SV40polyA-SV40promoter-ZeocinR-BGHpolyA-

hU6promoter	This study

pMaM519	pFA6a	GS_linker-mNeonGreen-SV40polyA-hU6promoter	This study

pMaM523	pFA6a	M1site(GS_linker)-mNeonGreen-SV40polyA-loxP-SV40promoter-ZeocinR-

BGHpolyA-hU6promoter-loxP-M2site	This study

pMaM526	pFA6a	M1site(GS_linker)-mNeonGreen-SV40polyA-loxP-hU6promoter-loxP-M2site	This study

pMaM530	pFA6a	M1site(GS_linker)-mNeonGreen(noATGs)-SV40polyA-loxP-hU6promoter-

loxP-M2site	This study

pMaM541	pFA6a	M1site(GS_linker)-mNeonGreen-SV40polyA-loxP-SV40promoter-PuromycinR-

BGHpolyA-hU6promoter-loxP-M2site	This study

pMaM549	pFA6a	M1site(GS_linker)-mNeonGreen(noATGs)-SV40polyA-loxP-SV40promoter-

ZeocinR-BGHpolyA-hU6promoter-loxP-M2site	This study

pMaM550	pFA6a	M1site(GS_linker)-mNeonGreen(noATGs)-SV40polyA-loxP-SV40promoter-

PuromycinR-BGHpolyA-hU6promoter-loxP-M2site	This study

pMaM552	pFA6a	M1site(GS_linker)-mScarlet-i(noATG)-SV40polyA-loxP-SV40promoter-

PuromycinR-BGHpolyA-hU6promoter-loxP-M2site	This study
